# Single-cell characterization of transcriptomic heterogeneity in lymphoblastoid cell lines

**DOI:** 10.1101/2020.09.24.311886

**Authors:** Elliott D. SoRelle, Joanne Dai, Jeffrey Y. Zhou, Stephanie N. Giamberardino, Jeffrey A. Bailey, Simon G. Gregory, Cliburn Chan, Micah A. Luftig

## Abstract

Lymphoblastoid Cell Lines (LCLs) are generated by transforming primary B cells with Epstein-Barr Virus (EBV) and are used extensively as model systems in viral oncology, immunology, and human genetics research. In this study, we characterized single-cell transcriptomic profiles of five LCLs and present a simple discrete-time simulation to explore the influence of stochasticity on LCL clonal evolution. Single-cell RNA sequencing revealed substantial phenotypic heterogeneity within and across LCLs with respect to immunoglobulin isotype; virus-modulated host pathways involved in survival, proliferation, and differentiation; viral replication state; and oxidative stress. This heterogeneity is likely attributable to intrinsic variance in primary B cells and host-pathogen dynamics. Stochastic simulations demonstrate that initial primary cell heterogeneity, random sampling, time in culture, and even mild differences in phenotype-specific fitness can contribute substantially to dynamic diversity in populations of nominally clonal cells.

## Introduction

Lymphoblastoid Cell Lines (LCLs) are immortalized cells prepared by *in vitro* transformation of resting primary B cells from peripheral blood with Epstein-Barr Virus (EBV).^1,2^ LCLs are used extensively in research as a model for EBV-associated malignancies including Diffuse Large B-Cell Lymphoma (DLBCL)^3,4^ and post-transplant lymphoproliferative disorder (PTLD).^5,6^ Because EBV is a non-mutagenic transformant in this context, LCLs constitute an important renewable source of human cells and genomic DNA that are used in immunological, genetic, and virology research.^7–11^

EBV is a double-stranded oncogenic gammaherpesvirus infecting over 90% of humans.^12^ *In vivo*, the virus typically establishes an asymptomatic persistent latent infection in episomal form^13,14^ within resting memory B cells.^15^ Latent infection can take one of several forms, each characterized by distinct programs of viral gene expression initiated from different promoters.^16^ For example, classical EBV infection within resting memory B cells *in vivo* is characterized by the Latency I program in which expression from the Q promoter yields a single viral protein, EBV Nuclear Antigen 1 (EBNA1), which functions to maintain the viral episome.^17^ Latency I, termed “true latency,” is established only after a complex progression of infection through pre-latency, Latency IIb, Latency III, and restricted forms of latency (e.g., Latency IIa), each occurring in distinct tissues within the body.^16^ While relatively infrequent, EBV can undergo lytic reactivation as a replication strategy.^18^

*In vitro*, the process of LCL production also necessarily involves multiple transitions in viral transcriptional programs. In the immediate-early stage of infection (the pre-latent phase), expression from the W promoter yields EBNA-LP, EBNA2, and several noncoding RNAs (EBERs, BHRF1 miRNAs, and BART miRNAs). A brief burst of lytic gene transcription (without lytic replication) is also observed during pre-latency.^19^ EBNA-LP and EBNA2 protein levels increase gradually within these early-infected cells, eventually leading to Latency IIb in which EBNA2 activation of the C promoter upregulates expression of EBNA1, EBNA3A-C, noncoding RNAs, and additional EBNA-LP and EBNA2.^20^ Latency IIb gene products induce hyperproliferation, a period of several days during which infected B cells divide every 10-12 hours.^21^ During hyperproliferation, EBNA1 mediates viral genome replication while EBNA3A-C inhibit host cell antiviral and tumor-suppression responses. Variance in virally-mediated rates of proliferation ensures that some infected cells undergo DNA damage-induced growth arrest^21,22^ while others continue to proliferate, eventually outgrowing as immortalized LCLs. LCLs largely exhibit the Latency III transcriptional profile, characterized by expression of all six EBNAs (EBNA-LP, EBNA1, EBNA2, and EBNAs 3A-3C) in addition to Latent Membrane Proteins 1 and 2 (LMP-1, LMP-2A/B) and noncoding RNAs.^23^ In Latency III, EBNA2 stimulates expression of LMP-1, a constitutively active Tumor Necrosis Factor Receptor (TNFR) homolog.^24^ LMP-1 signaling drives proliferation and survival via NFκB pathway activation,^25^ which has been shown to be essential for LCL outgrowth.^26^

Although studied extensively, complete characterization of the viral and host determinants of growth arrest versus immortalization of early-infected cells remains elusive.^27^ As one consequence, it is unclear whether or to what extent viral transformation may influence the resulting LCL cell populations. The possibility of significant phenotypic diversity within and across LCL samples warrants consideration, given the intrinsic variance of the human primary B cell repertoire^28,29^ and the multiplicity of viral transcription programs active in the journey to immortalization. Indeed, we recently described a gene expression program having low expression of LMP1 and NFκB targets which was unique to early infection (Latency IIb) relative to an otherwise identical population of LCLs.^30^ The wide distribution in LMP1 and NFκB target expression levels within an LCL has been characterized and ascribed to the dynamic sampling of a distribution of immune evasive states, at the fringes of which growth and survival can be compromised.^31–33^

In this study, we characterize the transcriptomic profiles of five different LCLs with single-cell resolution to assess inter- and intra-sample heterogeneity. Four of the sampled LCLs (two in-house and two commercial cell lines) were transformed with the prototypical B95-8 strain of EBV derived from an infectious mononucleosis patient,^34^ while a fifth sample (in-house) was prepared from cells transformed with the M81 strain isolated from a human nasopharyngeal carcinoma (NPC) sample.^35,36^ Primary cells used in establishing the five LCLs were isolated and transformed from a total of four donors; cells from one donor were transformed concomitantly to establish LCLs with each of the tested EBV strains. We observe B cell genetic heterogeneity in the form of differential heavy chain isotype expression across LCLs and, in three instances, within a sample. Further, comparable patterns of phenotypic variance with respect to NFκB pathway and plasma cell-like differentiation genes are evident in each LCL. Expression of host and viral genes indicate that individual cells within LCLs occupy a continuum of infection states. We also present an initial stochastic model to explore factors beyond the nuances of host-pathogen interactions that may generate profound phenotypic diversity within cultured cell lines. Our findings highlight some of the underappreciated complexity inherent within LCLs and broadly underscore the importance of understanding and accounting for sources of heterogeneity within presumptive cell lines.

## Results

### LCL generation and data provenance

Three LCLs were prepared in-house by infection of PBMCs from two donors (sample numbers 461 and 777) with one of two different EBV strains (B95-8 or M81). Each of these three samples (LCL 461 B95-8, LCL 777 B95-8, and LCL 777 M81) was prepared and processed using standard single-cell RNA sequencing workflows (see Experimental Methods). Two additional, publicly available datasets were obtained for commercially-available samples of the GM12878 and GM18502 LCLs, which were generated as previously reported by Osorio and colleagues.^37^ These five samples yielded single-cell RNA count matrices for subsequent analysis.

### LCL sample QC

Count matrices for the five samples exhibited similar feature, total RNA count, and mitochondrial gene distributions (**Figure 1 – figure supplements 1 & 2**) and were subjected to standardized QC thresholding (see Experimental Methods). Cell cycle marker expression (**Figure 1 – figure supplement 3**) was scored and regressed out during selection of highly variable genes as features to avoid clusters arising solely from cell cycle phase. Selected features were used to derive principal components which were evaluated (**Figure 1 – figure supplement 4**) and subsequently used for dimensional reduction (see Experimental Methods).

### Immunoglobulin isotype heterogeneity within and across LCL samples

The five LCL populations exhibit distinct immunoglobulin (Ig) profiles with respect to both gene expression levels and isotype frequencies (**Figure 1**). Three of the five samples (LCL 777 B95-8, LCL 777 M81, and GM12878) contain IgM^+^ and class-switched IgA^+^ and IgG^+^ subpopulations, whereas two samples (LCL 461 B95-8 and GM18502) almost exclusively expressed IgG (**Figure 1A**). Additionally, cells within each isotype class exhibit a wide range of Ig gene expression across all samples in an apparent class-independent fashion. No significant expression of IgE was observed in any of the five samples, consistent with the isotype’s rarity in the peripheral blood.^38,39^ Significant IgD transcript levels were observed in one sample (LCL 777 B95-8), where the gene’s expression was constrained to (and varied inversely with expression levels of) IgM^+^ cells (**Figure 1 – figure supplement 5**).

**Figure 1.**
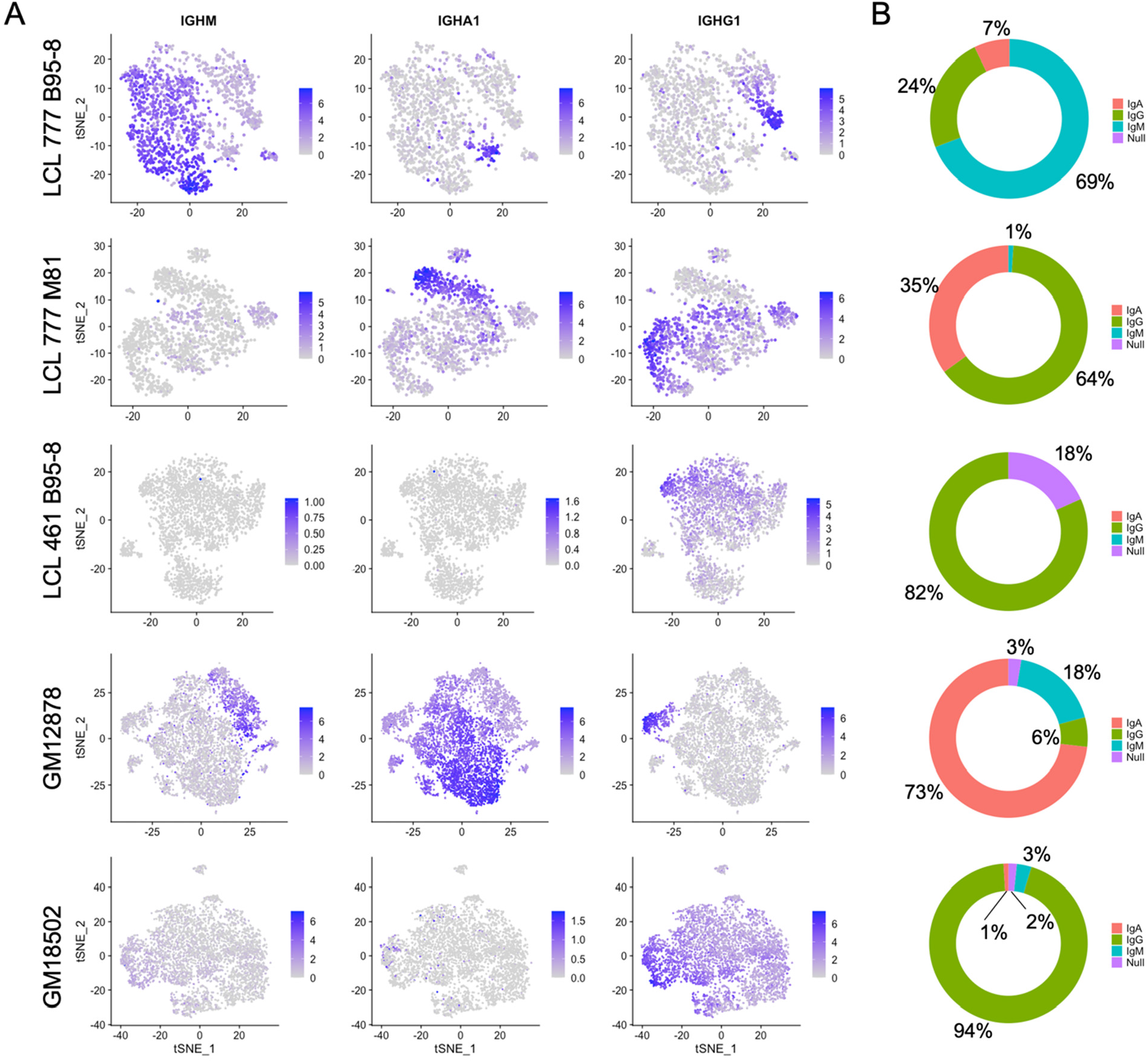
Immunoglobulin isotype heterogeneity within and across LCLs. **(A)** Relative expression of immunoglobulin heavy chain genes (IgM, IgA1, IgG1) in five LCLs analyzed by single-cell RNA sequencing. Data are represented by dimensional reduction (t-distributed Stochastic Neighbor Embedding) of principal components generated from feature selection following out-regression of cell cycle markers (see Experimental Methods). **(B)** Percent of cells in LCL population within each isotype class. Null classification represents cells exhibiting negligible immunoglobulin heavy chain expression.

The proportion of cells expressing each isotype varied substantially among LCLs (**Figure 1B**). IgG was the only isotype observed in LCL 461 B95-8. Cells in the GM18502 sample were also homogenous for IgG, although low levels of IgM transcripts are observed in up to half of the population. The proportion of cells () for IgM, IgA, IgG in LCL 777 B95-8 were (69%, 7%, 24%); in LCL 777 M81 were (1%, 35%, 64%); and in GM12878 were (6%, 73%, 18%). Abundance of Ig light chain gene (kappa or lambda) and heavy chain isoform expression are generally correlated with variable heavy chain expression in each of the five samples (**Figure 1 – figure supplements 5-14**). The isotype and clonal frequency differences between LCL 777 B95-8 and LCL 777 M81 are notable, given that these samples originated from the same donor and were transformed at the same time with different viral strains.

Differential Ig isotype expression is a significant source of variation in LCLs, as captured by the loadings from principal component analysis (PCA), typically within the first four PCs (**Figure 1 – figure supplement 15-19**). Consequently, differences in Ig isotype are effectively captured in dimensionally reduced datasets generated from PCs using t-distributed Stochastic Neighbor Embedding (tSNE) even at low clustering resolution. In samples with more homogenous isotype expression (LCL 461 B95-8 and GM18502), the relative Ig expression level is a significant factor in distinguishing clusters.

### Genes involved in B cell proliferation and differentiation exhibit inverse expression gradients

Across all samples, LCL populations display variable mRNA transcript levels for genes involved in cell proliferation, inhibition of apoptosis, response to oxidative stress, and differentiation (**Figure 2**). Gradients in Ig expression exhibit strong anticorrelation with expression of NFκB pathway transcripts (e.g., NFKB2, NFKBIA, EBI3) central to B cell proliferation and survival (**Figure 2A, Figure 2 – figure supplement 1A**). Similar gradients are observed for metabolic and oxidative stress response transcripts (e.g., TXN, PRDX1, PKM, LDHA, ENO1, HSP90AB1), however these transcripts are present more broadly (>80% of cells) and at higher levels than NFκB-related genes (20-30% of cells) in each sample (**Figure 2 – figure supplement 2**). While NFκB family gene expression is consistently anticorrelated with that of B cell differentiation factors, significant diversity exists in NFκB-high cells with respect to specific subunits including cREL, RELA, and RELB (**Figure 2 – figure supplement 3**). This implies differential intercellular NFkB dimer composition and, consequently, intra-sample variation in NFκB-mediated transcriptional programs. Expression of NFκB regulated BCL2 family members (e.g. BCL2L1/Bcl-xL and BCL2A1/BFL1) displays strong anticorrelation with Ig expression level. However, MCL1 and BCL2 mRNAs are more broadly expressed across cells within each LCL, while BCL2L2/BCL-W is only modestly expressed in LCLs (**Figure 2 – figure supplement 4**).

**Figure 2.**
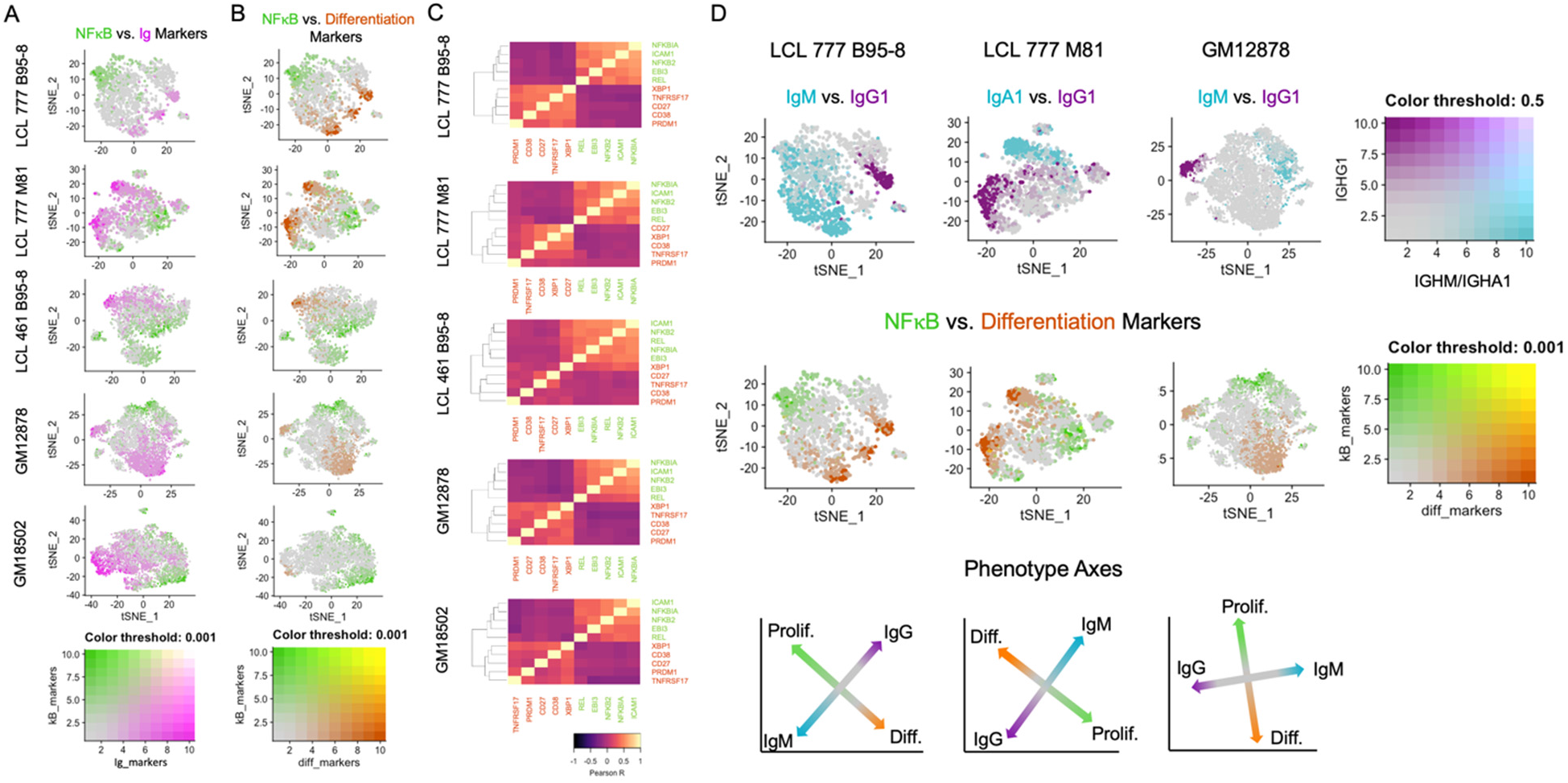
LCLs exhibit anticorrelated expression gradients of activation and differentiation genes. **(A)** Inverse expression gradients of immunoglobulin genes (IgM, IgA1, IgG1) in magenta and NFκB targets (NFKB2, NFKBIA, EBI3, ICAM1, BCL2A1) and TXN in green. **(B)** Similar inverse gradients of NFκB targets in green and B cell differentiation markers (TNFRSF17, XBP1, MZB1, CD27, CD38) in orange. **(C)** Pearson correlation maps and hierarchical clustering reveals negative correlation of differentiation (orange) and activation (green) gene sets and positive correlations between genes within each set. **(D)** In LCLs comprising multiple immunoglobulin isotypes, heavy chain class and differentiation/activation gradients constitute orthogonal (independent) axes of phenotypic variance.

Ig gradients are closely related to expression of differentiation and maturation markers (e.g., CD27, TNFRSF17/BCMA, XBP1, MZB1, PRDM1),^40–42^ which are likewise anti-correlated with NFκB pathway markers (**Figure 2B-C, Figure 2 – figure supplement 1B**). The apparent inverse relationship between these gene sets defines a major axis of phenotypic variance within LCL samples comprising multiple Ig isotypes (**Figure 2D**). The orthogonality of the pro-survival/differentiation and isotype class diversity axes implies that these two aspects of phenotypic variance are decoupled. Continuity between phenotypes resembling activated B cells (ABC) and antibody-secreting cells (ASC) is also captured in the expression profiles of key genes involved in the mutually antagonistic control of B cell state (**Figure 2 – figure supplement 5**).^43^ In this model, genes including PAX5 and IRF8 promote the ABC state; IRF4 and MKI67 (a G2M cell cycle marker) are markers of a transitional phenotype; and PRDM1 (BLIMP1) and XBP1 promote the ASC state. As cell cycle marker expression was regressed out, mitotic phase has negligible influence on the observed trends.

Whereas distinctions in Ig isotype class expression tend toward discrete partitioning, intra-isotype expression of differentiation and maturation genes reflects a continuum of transcriptomic states and cellular functions. Thus, within a given isotype, elevated Ig heavy chain expression is negatively correlated with proliferation/anti-apoptotic gene expression and positively correlated with maturation/differentiation gene expression. These relationships are most readily evident in LCL samples consisting of a single class-switched population, such as GM18502 (**Figure 2 – figure supplement 1C**).

Finally, the viral EBNA2 and EBNA3 proteins are responsible for transcriptional regulation that we specifically interrogated within the single cell data. The direct EBNA2 targets RUNX3 and FCER2/CD23 correlated with NFκB expression (**Figure 2 – figure supplement 6**).^44^ Indeed, the expression of RUNX3 and FCER2/CD23 was anticorrelated with Ig expression consistent with the known role of EBNA2 in suppressing IgH transcription.^45^ In contrast, the EBNA3 repressed targets including CXCL9, CXCL10, BCL2L11/BIM and ADAMDEC1 were uniformly repressed (**Figure 2 – figure supplement 7**) consistent with the role of histone and DNA methylation in maintain gene repression of EBNA3 targets.^46–48^

### Viral state heterogeneity affects host expression profile distributions in LCLs

Clusters with high EBV lytic gene expression are observed in two of the three datasets (LCL 777 B95-8 and LCL 777 M81) aligned against the human reference genome containing the viral genome as an extra chromosome (see Experimental Methods) (**Figure 3**). Lytic cluster cells are small, accounting for 2.2% and 0.9% of the LCL 777 B95-8 and LCL 777 M81 cell populations, respectively (**Figure 3A**). The higher rate of lytic cell capture in the B95-8 sample relative to the M81 sample is somewhat surprising, as the M81 strain is known for increased frequency of lytic reactivation; however, this disparity may originate from the nature of single-cell sample preparation method (see Discussion).^49^

**Figure 3.**
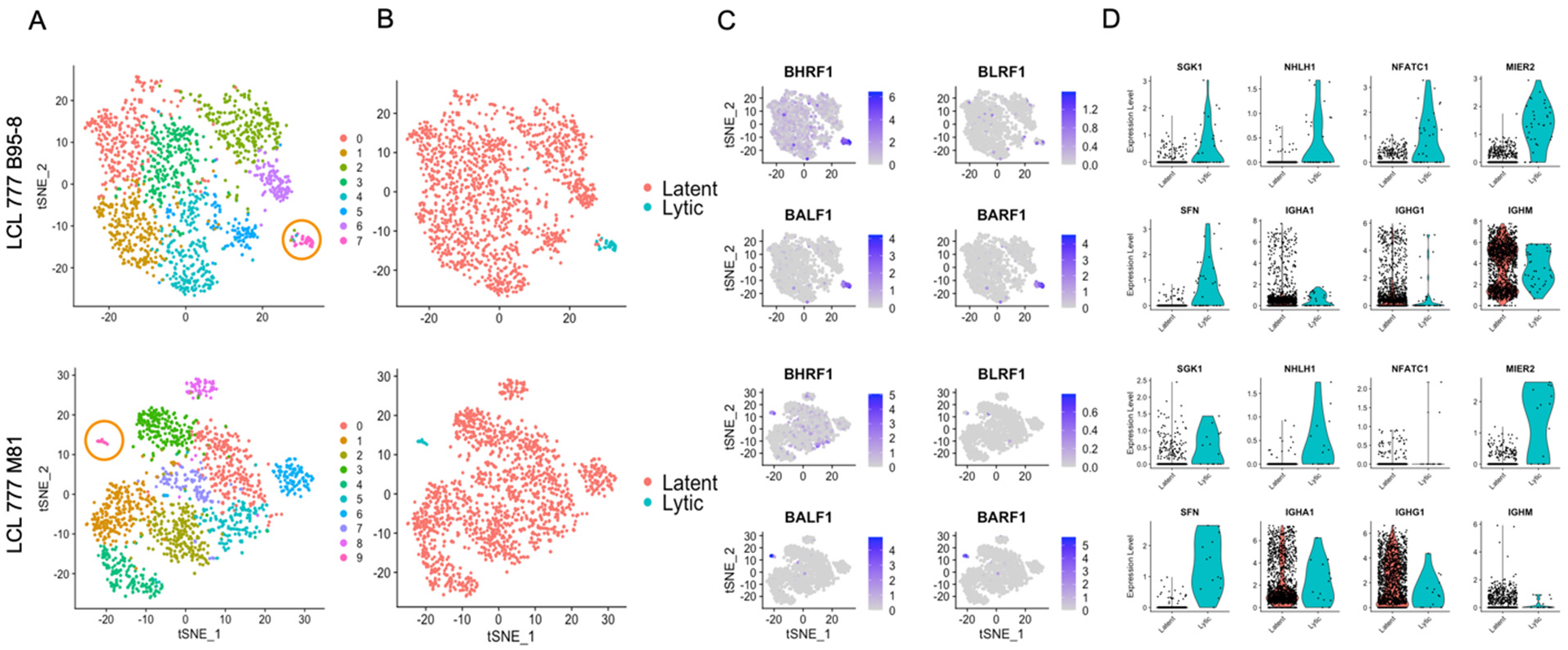
Viral and host gene expression in lytic cell subpopulations. **(A)** Clustering of dimensionally reduced datasets for LCL 777 B95-8 and LCL 777 M81. **(B)** Grouping of cell clusters into latent (red) and lytic (cyan) cells based on viral and host gene expression signatures of principal components. **(C)** Relative expression of four representative EBV lytic genes (BHRF1, BLRF1, BALF1, and BARF1) is elevated in lytic cell subpopulations. **(D)** Lytic cell clusters exhibit elevated expression of several host cell genes (SGK1, NHLH1, NFATC1, MIER2, SFN) relative to latently infected cells. While under-sampled due to subpopulation size, immunoglobulin class frequencies in lytic cells roughly reflect the population-wide frequencies.

The presence or absence of viral lytic transcripts is a significant source of phenotypic variance in these samples, as reflected in population groupings by viral state (**Figure 3B**) and principal component loadings (**Figure 1 – figure supplements 15 & 16**, PC_3 and PC_7, respectively). Lytic cells can be identified confidently from high expression of EBV genes including BLRF1, BALF1, and BARF1, among others (**Figure 3C**). BHRF1 expression is also elevated in lytic cells, although BHRF1 transcripts are ubiquitous at low levels sample wide. This is likely because BHRF1 can be expressed during both latent and lytic phases of EBV infection from different promoters.^50^

While the absolute number of lytic cells in each sample is low, the data indicate that the lytic cells are polyclonal with respect to Ig heavy chain expression, display upregulation of several host genes including NFATC1, MIER2, SFN, and SGK1, and exhibit heterogeneous NFκB expression (**Figures 3D, Figure 1 – figure supplements 5 & 6**). Ig isotype distributions in lytic cell clusters appear roughly proportional to the whole-sample distributions. NFATC1, MIER2, SFN, and SGK1 transcript levels were queried for GM12878 and GM18502 samples to test whether the presence of lytic cell subpopulations might be inferred from host gene expression. A sub-cluster representing a small percentage of cells in GM12878 (<0.5%) were found to co-express MIER2 and NFATC1. Negligible expression of either gene was observed in GM18502 (**Figure 1 – figure supplements 8 & 9**).

### Loss of mitochondrial and Ig expression in subpopulations under oxidative stress

Three of the five samples (LCL 461 B95-8, GM12878, and GM18502) contain clusters that exhibit metabolic transcriptional profiles in stark contrast with typical expression in each population (**Figure 4**). Cells within these clusters account for 1-4% of the three samples after QC (**Figure 4A**) and are most notable for their low expression of mitochondrial genes (**Figure 4B**). In the case of LCL 461 B95-8 and GM18502, these cells are the first to partition from the rest of the sample at low clustering resolution (**Figure 4 – figure supplements 1-5**).

**Figure 4.**
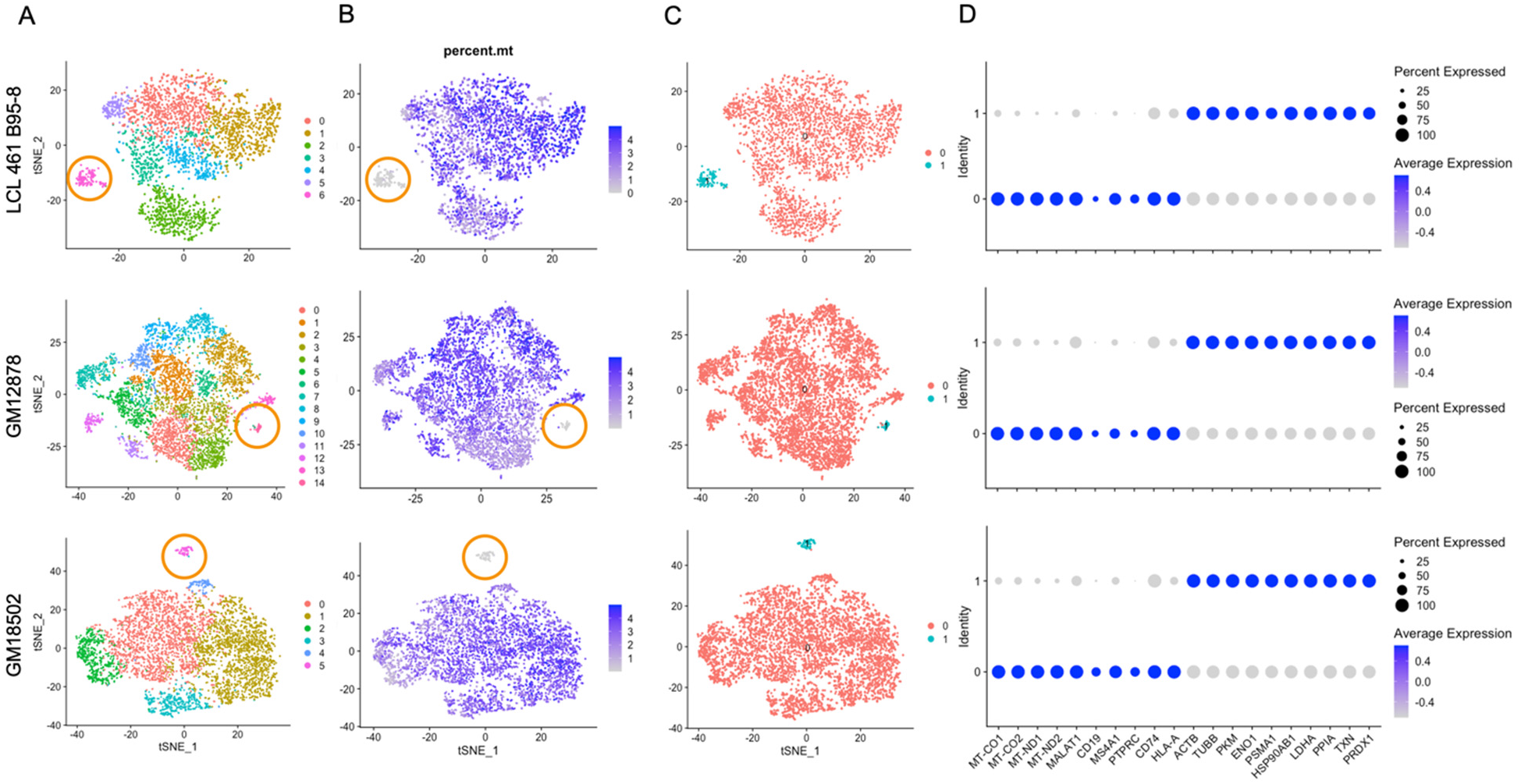
LCL subpopulations exhibiting reduced mitochondrial gene expression and elevated metabolic and oxidative stress genes. **(A)** Clustering of dimensionally reduced datasets for LCL 461 B95-8, GM12878, and GM18502. **(B)** Distinct clusters within each of these samples are defined by uncharacteristically low mitochondrial gene expression. **(C)** Grouping of cell clusters to partition “mito-low” cells (cyan) for differential expression comparison. **(D)** Mito-low cells exhibit reduced expression of cytochrome oxidase (MT-CO1, MT-CO2), NADH-ubiquinone oxidoreductase (MT-ND1, MT-ND2), MALAT1, and numerous lymphoid and B-cell lineage markers (CD19, MS4A1/CD20, PTPRC/CD45, CD74, HLA-A). Mito-low cells exhibit increased expression of genes associated with cytoskeletal rearrangements (ACTB, TUBB), metabolic stress (PKM, ENO1, LDHA) protein folding/degradation (HSP90AB1, PSMA1, PPIA), and oxidative stress (TXN, PRDX1).

Compared to the rest of each sample, these atypical cells exhibit significantly depleted levels of cytochrome c oxidase (MT-CO, complex IV) and NADH-ubiquinone oxidoreductase subunits (MT-ND, complex I) as well as a lack of canonical markers of lymphoid (e.g., PTPRC [CD45], CD74), B cell-specific lineage (e.g., CD19, MS4A1 [CD20]), and in some cases, MHC class I & II antigen presentation (e.g., HLA-A,B,C, HLA-DR) (**Figures 4C-D, Figure 1 – figure supplements 7 & 9, Figure 4 – figure supplement 6**).

Expression of genes involved in oxidative stress (TXN, PRDX1), unfolded protein responses (PPIA, HSP90AB1), metabolic shunt pathways (PKM, ENO1, LDHA), and cytoskeletal rearrangements (ACTB, TUBB) is enriched consistently in this subset relative to the bulk population in each of the three LCLs (**Figure 4D, Figure 2 – figure supplement 2**). Ig heavy chain transcripts are notably absent from these subpopulations, although some degree of light-chain expression is observed (**Figure 1 – figure supplements 7-9**). While these cells are on the low end of the population distribution with respect to total RNA counts and unique feature RNAs (**Figure 4 – figure supplement 7**), the measured values are consistent with intact, viable cells.

### A stochastic model for LCL phenotypic heterogeneity

A simple stochastic simulation based on a discrete-time Markov chain model^51^ was developed to understand better the factors that may influence phenotypic heterogeneity observed in LCLs, using Ig isotype frequencies as an example (**Figure 5**). In principle, the simulation may be adapted to any set of phenotypes within a sample. For additional details regarding model parameters and assumptions, please see the Experimental Methods (Stochastic Simulations) and refer to the source code (supplementary file: “ig_evo_sim.py”).

**Figure 5.**
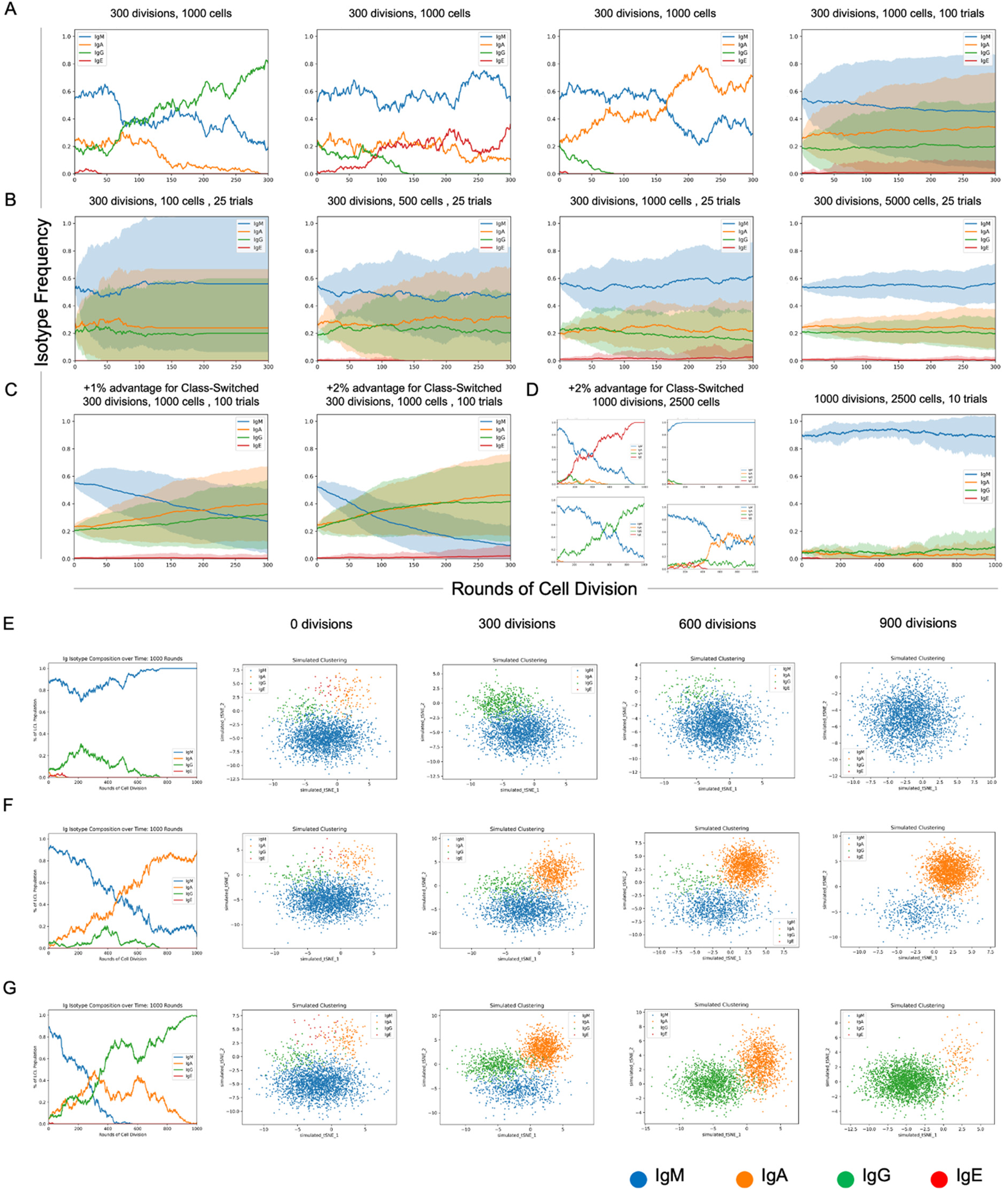
Stochastic simulation of heterogeneous LCL evolution. **(A)** Stochastic immunoglobulin isotype frequency evolution. Three random single-trial simulations initiated from the same starting class frequencies are presented, assuming equal likelihood of proliferation across isotype classes (n = 1000 cells). The last panel shows mean and standard deviation for outcomes from 100 trials simulated from the same parameters. **(B)** Simulation of a founder effect. Population under-sampling (modeled by comparing results from 25 trials using n = 100, 500, 1000, and 5000 cells, left-to-right panels) increases outcome variance and accelerates convergence to a single isotype. **(C)** Effect of phenotype-specific fitness advantages. Simulation results are presented for scenarios in which class-switched isotypes (IgA, IgG, IgE) have a 1% (left panel) or 2% (right panel) fitness advantage over IgM cells. **(D)** Four random single-trial simulations over long periods of time (1000 division rounds) with a 1% fitness advantage for class-switched cells (left panels) compared to 10 trials over the same period with equal fitness across classes. **(E)** Single-trial isotype frequency evolution and corresponding simulated clustering (see Experimental Methods) in the case of equal proliferation probability. Starting frequencies of IgM, IgA, IgG, IgE cells are 89%, 5%, 5%, and 1%, respectively. **(F)** As in E, with a 1% fitness advantage for class-switched cells. **(G)** As in E, with a 2% advantage for class-switched cells.

In the present implementation, changes in Ig isotype frequency can be simulated in discrete steps (rounds of cell division) as a function of initial phenotype frequencies, population sampling (with replacement), and potential differences in phenotypic fitness captured as fixed, (un)equal isotype-specific proliferation probabilities. The model assumes a fixed cell death rate across all isotypes in any given division round. The number of simulated trials can be adjusted to capture individual stochastic realizations or probabilistic outcome distributions. Each parameter and assumption can be adjusted by the user for tailored applications.

Three randomly selected realizations and averaged outcomes (trials = 100) of the model for a fixed sample size (n = 1000 cells) demonstrate the effects of intrinsic stochasticity on the evolution of phenotype proportions over many rounds of cell division (rounds = 300), even when each phenotype confers equivalent fitness (**Figure 5A**). In the case of equal fitness and sufficient sample size, initial phenotype frequencies are a key determinant of whether the most prevalent phenotype will change over time as a result of stochasticity.

The effect of sample size on inter-trial variance can be substantial, even when cell populations are sampled with replacement to maintain phenotype proportions in each round (**Figure 5B**). Mean phenotype proportions are generally conserved, whereas trial standard deviation decreases as the sample size increases (trials = 25, rounds = 300, n = 100, 500, 1000, or 5000 cells). This is generally expected, since undersampling increases the likelihood that phenotype frequencies in the drawn sample will deviate from those of the population, even in the case of replacement.

It is notable that minor differences in relative fitness (1-2%) can lead to dramatic changes in isotype distributions over time (**Figure 5C**). The rate of such change is proportional to the magnitude(s) of fitness differences (n = 1000 cells, rounds = 300). Four randomly selected clonal evolution trajectories realized with a modest fitness advantage (2%) for class-switched cells (IgA, IgE, and IgG) reveal the potential for drastic variations when multiple rare phenotypes with a fitness advantage exist (n = 2500 cells, trials = 10, rounds = 1000). Thus, rare cells may become prevalent or even dominant over time if they exhibit only slightly greater fitness relative to other cells in some environmental context (e.g., cell culture). In such cases, observed phenotype frequencies can deviate wildly from expectations of equal fitness over time (**Figure 5D**).

Cluster simulation was implemented by random sampling from four arbitrary, isotype-specific 2D normal distributions based on empirical observations that Ig isotypes yield distinct clusters in dimensionally reduced single-cell RNA-seq data (**Figures 5E-G**). Simulated clusters were generated from randomly selected trials initiated from the same initial phenotype distribution (IgM = 89%; IgA = 5%; IgG = 5%; IgE = 1%) at three different relative fitness advantages (0%, 1%, and 2%) for class-switched isotypes. In all cases, the proportion of observed cells in each cluster fluctuates over time. As expected, the presence or absence of observed phenotypic heterogeneity (in this example, isotype polyclonality) in a cell population is a complex function of relative frequency, fitness, sampling (i.e., bottlenecks), stochasticity, and time.^52,53^

## Discussion

### Ig isotype heterogeneity in LCLs

LCL clonality is known to change over time, although the factors involved in this evolution are not fully characterized.^54^ PBMC derivation from multiple donors is an obvious source of cellular heterogeneity in the analyzed samples presented herein. B cells from peripheral blood (≈ 5-10% of all lymphocytes) comprise wide ranges of naïve (≈ 50-80%, mean ≈ 65%) and memory (≈ 15-45%, mean ≈ 30%) cells, with immature/transitional and plasmablasts accounting for smaller proportions (≈ 1-10%, mean ≈ 5% and ≈ 0.5-4.5%, mean ≈ 2%, respectively).^29^ Within the memory cell compartment, proportions of non-switched (IgM) and switched memory (IgA, IgG, and technically, IgE) are also likely donor-specific. The negligible number of IgE^+^ cells present across the samples can be explained by the isotype’s low frequency in the peripheral blood.^38^

It is evident from LCL 777 B95-8 and LCL 777 M81 samples that inter-donor differences cannot fully explain the observed isotype heterogeneity in LCLs. While it may be tempting to attribute the observe differences to infection with different viral strains, there is ample experimental evidence that EBV infection does not induce class-switching.^55^ The disparity in isotype frequencies is notable since these samples were transformed, cultured, prepared, and sequenced in parallel (i.e., under equivalent conditions and within the same interval).

The polyclonality exhibited within LCL 777 B95-8 and LCL 777 M81 contrast with the dominance of a single isotype in LCL 461 B95-8 and GM18502 samples (in each case, IgG). The only notable difference between LCL 461 B95-8 and LCL 777 B95-8 is that the former sample was in culture substantially longer prior to single-cell library preparation. Given that the GM18502 line was derived more than a decade ago, these observations implicate the influence of culture period in altering significantly the isotype proportions present within LCLs, which is altogether consistent with known (and profound) challenges associated with cell culture.^56–58^ In this regard, the data from GM12878 merit remark. The finding of polyclonality in this sample is surprising, given that GM12878 has been in culture over a timescale comparable to GM18502.^59^ Forgoing the possibility of errors in sample handling or procurement, the persistence of genetic heterogeneity in this line is both intriguing and potentially confounding. Whether or to what extent cellular diversity may influence observed results will inevitably vary on a study-specific basis, but the possibility of sample-intrinsic variance should be considered even when homogeneity is presumed.^8,60^

Multiple isotypes within an LCL sample guarantee clonal diversity, but the presence of a single isotype does not necessarily ensure the inverse (intra-sample homogeneity). While not in the scope of the present study, B Cell Receptor (BCR) 5’ single cell sequencing of LCL samples could provide insight into variable regions and as to whether subpopulations of a given isotype are the progeny of one or multiple founder cells (and whether the answer to this question changes over time).

### Viral origins of LCL phenotypic variance

NFκB pathway signaling is constitutively activated by viral Latent Membrane Protein 1 (LMP1) in EBV-transformed B cells.^25^ LMP1-induction of the NFκB pathway is necessary for LCL survival;^26,61,62^ however, the observed intra- and inter-LCL variance in transcript levels of NFκB and several of its transcriptional targets add nuance to this picture. Similar profiles of NFκB pathway transcript levels across samples may constitute a snapshot of the most probable distribution arising from stochastic NFκB target expression induced by EBV infection. This may arise from a transcriptional bursting mechanism in which mRNA transcript levels in each cell fluctuate over time (as a Poisson process) while the proportion of cells containing *n* transcripts in a population at any given time is roughly constant.^63–67^ Alternatively, or perhaps additionally, variation in NFκB pathway activity may be a manifestation of the different viral latency states present within each sample, as indicated by correlation with host markers of latency IIb and III.

The distinct anti-correlation between NFκB/viral latency program and B lymphocyte differentiation genes is noteworthy. While a mechanism imparting causality to this relationship is not yet fully clear, recent time-resolved bulk transcriptomic data revealed that EBV-induced plasma cell phenotypes (including upregulation of Xbp1) developed as early as the pre-latent phase of infection (1-14 days).^27^ Correlated expression of MZB1 with XBP1, TNFRSF17, CD27, and CD38 support the model that the development of plasma cell characteristics is reminiscent of germinal center differentiation. Single-cell data adds complexity to this finding and its consequences for LCL heterogeneity even after long-term outgrowth. Specifically, EBV transformation *in vitro* appears to maintain B cells along a continuum of differentiation states, each with varying degrees of similarity to phenotypes observed *in vivo*.^16^ In the case of LCL generation, the multiple transcriptional programs of the transformant likely constitute an inescapable source of phenotypic heterogeneity.

The low number of observed lytic cells is likely a consequence of EBV’s predominant latency and the fact that lytic reactivation is by nature somewhat incompatible with single-cell RNA-seq methods. However, these small subpopulations provide an interesting case for examination. The spatial proximity of lytic clusters in LCL 777 B95-8 and LCL 777 M81 to plasma-like clusters resulting from tSNE dimensional reduction implied phenotypic similarity, however we found that this is likely an artifact of the tSNE algorithm since UMAP dimensional reduction did not preserve this proximity. Notwithstanding, XBP1 upregulation in plasma cells has been shown to transactivate the viral Z promoter and induce lytic reactivation.^68,69^ Lytic cells also display relatively high and polyclonal Ig heavy chain expression in addition to other shared characteristics with plasma-like cells (reduced expression of NFκB and its targets). By contrast, lytic cells exhibit notably reduced levels of B cell differentiation transcripts. Thus, viral transcription changes in dynamic response to host cell programs (and vice versa) contribute to the observed LCL diversity. Prior work has shown that the viral proteins EBNA3A and EBNA3C suppress plasma-like phenotypes during EBV latency establishment.^70^ The possibility that EBV may undergo lytic reactivation in response to plasma cell differentiation as a means of maintaining persistent latent infection is a topic of future interest.

Host genes upregulated within lytic cluster cells (e.g., NFATC1, MIER2, SFN, SGK1) represent a limited subset of transcription factors associated with B (and T) lymphocyte activation^71,72^, several of which have been recently identified at various degrees of enrichment within lytic cells.^73^ The presence of NFATC1 is particularly notable considering the recent report of this factor contributing to the spontaneous lytic phenotype of type 2 EBV by upregulating expression of BZLF1 to promote the lytic gene expression cascade.^74^

Although PC loadings reveal substantial upregulation of more than a dozen EBV lytic genes, cells within the lytic clusters curiously lack expression of BZLF1, which plays a role in the latent-to-lytic transition.^18^ The absence of BZLF1 reads (and low mRNA counts generally) ostensibly may result from factors including naturally low transcript abundance, reduced transcript capture efficiency, and/or reduced efficiency of reverse transcription to cDNA owing to RNA secondary structural motifs.^75^

### “Marker-less” subpopulations

The small populations of cells in LCL 461 B95-8, GM12878, and GM18502 characterized by low mitochondrial gene expression and a dearth of canonical B cell markers are curiosities. These cells share similarities with exhausted plasma cells, most notably an apparent loss of Ig heavy chain expression while retaining moderate kappa and light chain expression,^76,77^ and hallmarks of oxidative stress including upregulated thioredoxin expression.^78–81^ Low levels of NFκB pathway transcripts in these clusters most closely resemble expression profiles of cells with a plasma-like phenotype in the same samples. It is unlikely that these cells are immature, naïve, or transitional B cells, given that neither IgM nor IgD expression are observed. Loss of lineage marker expression is suggestive of a tumor-like phenotype.^82^

### Factors in the evolution of subclonal heterogeneity

Cellular diversity abounds even within presumptive clonal lines. For LCLs generated from EBV-transformed primary B cells, the list of parameters affecting the cell population’s phenotypic profile includes donor-specific frequencies of non-switched and switched memory B cells, heterogeneous states of viral infection, phenotype-specific differential fitness in culture, stochasticity, and time. By definition, some degree of differential fitness exists among cells in each sample as a consequence of the variability in pro-survival, proliferation, and anti-apoptotic genes. However, as a principle of evolution, phenotypic differences do not necessarily have to be selected directly; they may simply be carried over in cells possessing other selected features. With respect to the stochastic model presented herein, the simulated phenotype advantage of class-switched memory vs. non-switched memory cells need not be construed as originating from heavy chain isotype expression.

Experimental procedures including cell passaging and the initial transformation itself may contribute to variance among LCLs. As an illustration, consider that 1 million PBMC has around 25,000 B cells, of which 7,500 (30%) on average are memory cells of various classes. If the rate of transformation leading to LCL outgrowth is 10%, then ≈ 750 memory cells out of 1 million PBMCs define the initial isotype frequency of the eventual LCL. This sample size is small relative to the donor’s total memory B cell compartment and may lead to founder cell effects. Consequently, B cell population undersampling may be a foregone conclusion in the context of LCL preparation.

## Conclusion

Single-cell RNA sequencing reveals that LCLs including widely used commercial lines exhibit substantial phenotypic diversity. During the early stages of LCL generation, EBV infection drives cell proliferation by mimicking the process of B cell activation. After successful LCL outgrowth, infected B cells occupy a range of phenotypic states along a continuum between activation and plasma cell differentiation and, in some cases, exhibit signs of lytic reactivation. The diversity observed within LCLs (and cultured lines generally) can originate from intrinsic heterogeneity within primary cells, transcriptional programs of the viral transformant, and the realization of inherently stochastic processes (including certain gene expression programs) over time. The data reported herein enable extensive hypothesis generation and interrogation of aspects of B cell biology, EBV pathogenesis, and host-virus interactions. Moreover, this work highlights the importance of considering the possible sources and experimental consequences of cell population heterogeneity when using cultured cell lines.

## Acknowledgments

We would like to acknowledge the assistance of the Duke Molecular Physiology Institute Molecular Genomics core for the generation of the data for the manuscript.

## Experimental Methods

### PBMC isolation and transformation with EBV

Whole blood samples from two normal donors (777 and 461) were obtained from the Gulf Coast Regional Blood Center. PBMCs were isolated from each sample by Ficoll gradient (Sigma, # H8889). CD19^+^ B cells were extracted from each PBMC sample through magnetic separation (BD iMag Negative Isolation Kit, BD, # 558007). Purified B cells were cultured in RPMI 1640 media supplemented with 15% fetal calf serum (FCS, vol./vol., Corning), 2 mM L-glutamine, penicillin (100 units/mL), streptomycin (100 μg/mL, Invitrogen), and cyclosporine A (0.5 μg/mL).

B95-8 and M81 strains of EBV were generated from the B95-8 Z-HT and M81 cell lines, respectively, as described previously.^83^ Separate bulk infections of B cells were performed by incubating donor B cells with B95-8 Z-HT or M81 supernatants for 1 h at 37°C, 5% CO_2_ to produce the following cultures: 777_B95-8, 777_M81, and 461_B95-8. After virus incubation, cells were rinsed in 1x PBS and resuspended in R15 media. LCL outgrowth was achieved from each of these three samples, resulting in LCL_777_B95-8, LCL_777_M81, and LCL_461_B95-8.

### Cell culture

All three in-house LCL samples were cultured in supplemented RPMI media as described above, substituting 10% FCS instead of 15% FCS. Prior to single-cell sample preparation, LCL_777_B95-8 and LCL_777_M81 were maintained in culture for approximately one month, whereas LCL_461_B95-8 was cultured for longer than six months. Immediately prior to single-cell sample preparation, LCLs were resuspended and disaggregated.

### LCL samples and data

LCL_777_B95-8, LCL_777_M81, and LCL_461_B95-8 were created as described above. LCLs GM_12878 and GM_18502 were obtained, prepared, sequenced, and aligned as described by Osorio and colleagues.^37^ Briefly, these samples were obtained from the Coriell Institute for Medical Research, cultured for several days, then prepared as single-cell GEMs (Gel bead in Emulsions) with the 10x Genomics Chromium system using version 2 chemistry for total RNA. Single-cell sequencing libraries were generated using established 10x Genomics protocols, and sequencing was performed with a Novaseq 6000 (Illumina, San Diego). Unique Molecular Identifier (UMI) count matrices were generated from these samples by using CellRanger v.2.1.0 with alignment to the hg38 version of the human reference genome. Additional information about the experimental handling and acquisition of data for GM12878 and GM18502 is provided in the original reference.^37^ Gene-barcode matrix files for each sample were downloaded from the Gene Expression Omnibus (accession ID: GSE126321) and subsequently analyzed along with data from LCL_777_B95-8, LCL_777_M81, and LCL_461_B95-8 samples, while the LCL_461_B95-8 sample was run in a separate experimental batch.

### Single-cell RNA sample preparation, and sequencing

Single-cell RNA samples for LCL_777_B95-8, LCL_777_M81, and LCL_461_B95-8 were prepared using the General Sample Preparation demonstrated protocol from 10x Genomics (10x, Manual Part # CG00053) adapted from the original published methods.^49^ Briefly, disaggregated LCLs were resuspended in fresh 1x PBS supplemented with 0.04% BSA, stained with trypan blue to assess viability, and counted using a hemocytometer for preparation to target concentration.

Single-cell libraries for sequencing were prepared from each sample using the methods described in the 10x Genomics Single Cell 3’ Reagent Kit Protocol (v2 chemistry, Manual Part # CG00052). In brief, GEMs were prepared using the 10x Chromium Controller, after which cDNA synthesis and feature barcoding were performed and sequencing libraries for each sample were constructed. Sequencing runs were performed on an Illumina HiSeq 3000/4000 (Illumina, San Diego). Samples for LCL_777_B95-8 and LCL_777_M81 were sequenced in a pooled run in a single HiSeq lane.

Raw base call files (*bcl.gz) from sequencing runs were processed using CellRanger v.2.0.0 to generate fastq files (*fastq.gz) via CellRanger’s ‘mkfastq’ command. CellRanger’s ‘count’ command was then used to align reads from the three in-house LCL samples to the human reference genome (hg38) with the Type 1 EBV reference genome (NC_007605) concatenated as an extra chromosome (reflecting the episomal nature of the EBV genome within infected B cells). This process yielded gene-barcode matrices (UMI count matrices) for subsequent analysis.

### Sample QC, analysis, and visualization

UMI count matrices for all five LCL samples were analyzed using the Seurat single-cell analysis package for R (Seurat v.3.1.5).^84,85^ Filtered barcode matrices were loaded into Seurat, after which genes present in fewer than three cells and cells expressing fewer than 200 unique RNA molecules (features) and more than 65000 unique features were filtered out. Additionally, cells in which mitochondrial genes accounted for greater than 5% of all transcripts were excluded from analysis. Beyond the uniform application of QC steps, we did not investigate the potential for batch-specific effects across the five samples run in four experiments. After QC thresholding, feature data were normalized and scored for cell cycle markers. Cell cycle scoring was used to regress out S and G2M gene features to remove variance (and unwanted effects on clustering) in the datasets arising from cell cycle phase. Cell cycle-corrected data were then scaled, and selection was performed to find the highest-variance features. Principal Component Analysis (PCA) was performed on selected (n = 2000) variable features, and PCs were subsequently used to define distinct subpopulations within each of the five samples. For visualization, PCs were used to generate clusters at various resolutions and dimensionally reduced using t-distributed Stochastic Neighbor Embedding (tSNE). The R code used to process data and produce figures presented in this manuscript is provided as a supporting file (“ig_evo_sim.py”) and is available on GitHub (https://github.com/esorelle/ig-evo-sim).

### Stochastic simulations

The concept of a discrete-time Markov chain was adapted to simulate the evolution of phenotype frequencies, using immunoglobulin heavy chain isotype distributions within LCLs as an example. Briefly, the simulation takes as input a cell population of size n comprising B cells of different Ig heavy chain isotype classes at user-defined initial frequencies, fixed probabilities of proliferation in synchronous rounds of cell division, and a constant cell death rate assumption (also user-defined). Within the scope of computational feasibility, users can specify the number of rounds of cell division to simulate and the number of simulation trials to run. Additionally, users may choose to generate simulated cluster data modeled from distinct 2D normal distributions for each isotype for a specified number of trials at fixed intervals (i.e., every n^th^ cell division round). The simulation was implemented in Python, and the code used to generate the simulated data is provided as a supporting file. The code is also available at (add as public GitHub repo) and may be freely implemented and modified.

## Author Information

**Elliott D. SoRelle**

**Contributions:** Conceptualization, Data curation, Formal analysis, Investigation, Validation, Writing – original draft, Writing – review and editing

**Competing Interests:** No competing interests declared

**Joanne Dai**

**Contributions:** Conceptualization, Data curation, Formal analysis, Investigation, Methodology, Validation, Writing – reviewing and editing

**Competing Interests:** No competing interests declared

**Jeffrey Zhou**

**Contributions:** Data curation, Formal analysis, Investigation, Validation

**Competing Interests:** No competing interests declared

**Stephanie Giamberardino**

**Contributions:** Data curation, Formal analysis, Investigation, Methodology, Writing – reviewing and editing

**Competing Interests:** No competing interests declared

**Jeffrey A. Bailey**

**Contributions:** Supervision, Writing – reviewing and editing

**Competing Interests:** No competing interests declared

**Simon Gregory**

**Contributions:** Formal analysis, Methodology, Supervision

**Competing Interests:** No competing interests declared

**Cliburn Chan**

**Contributions:** Supervision, Writing – reviewing and editing

**Competing Interests:** No competing interests declared

**Micah A. Luftig**

**Contributions:** Conceptualization, Formal analysis, Funding acquisition, Supervision, Writing – reviewing and editing

**Competing Interests:** No competing interests declared

## Source Data Files

Raw sequencing data for the three previously unpublished samples (LCL_777_B958, LCL_777_M81, and LCL_461_B958) are deposited in the NCBI Sequence Read Archive (SRA) and can be accessed along with processed data from the NCBI Gene Expression Omnibus (GEO, Series Accession: GSE158275).

## Source Code

R code used for UMI count matrix processing, analysis, and figure generation is provided as a supplementary file. Python code used for clonal evolution simulations is available on github (https://github.com/esorelle/ig-evo-sim).

## Figure Supplements

**Figure 1 – figure supplement 1.**
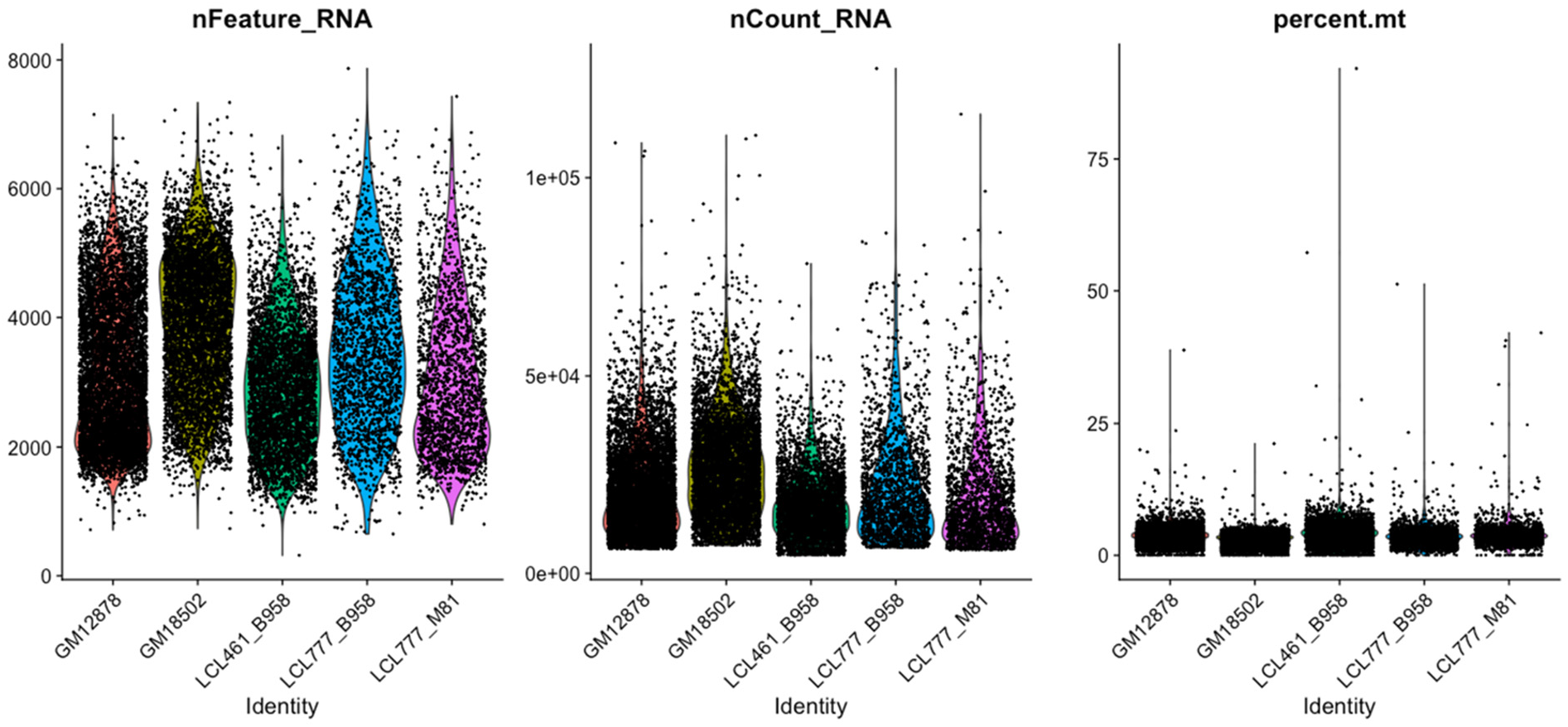
Distributions of features used for QC across five LCL samples.

**Figure 1 – figure supplement 2.**
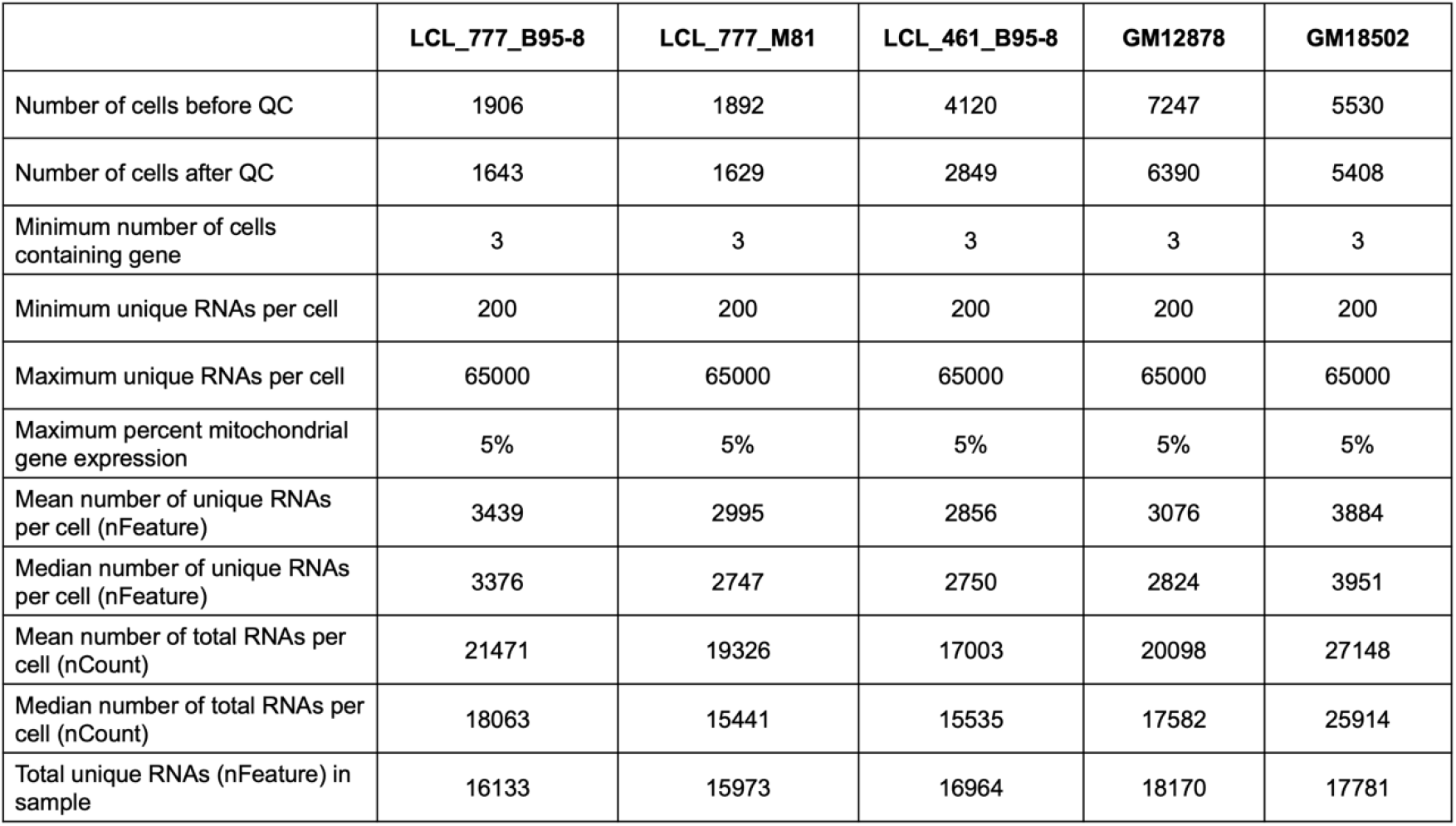
Summary of QC statistics across five LCL samples.

**Figure 1 – figure supplement 3.**
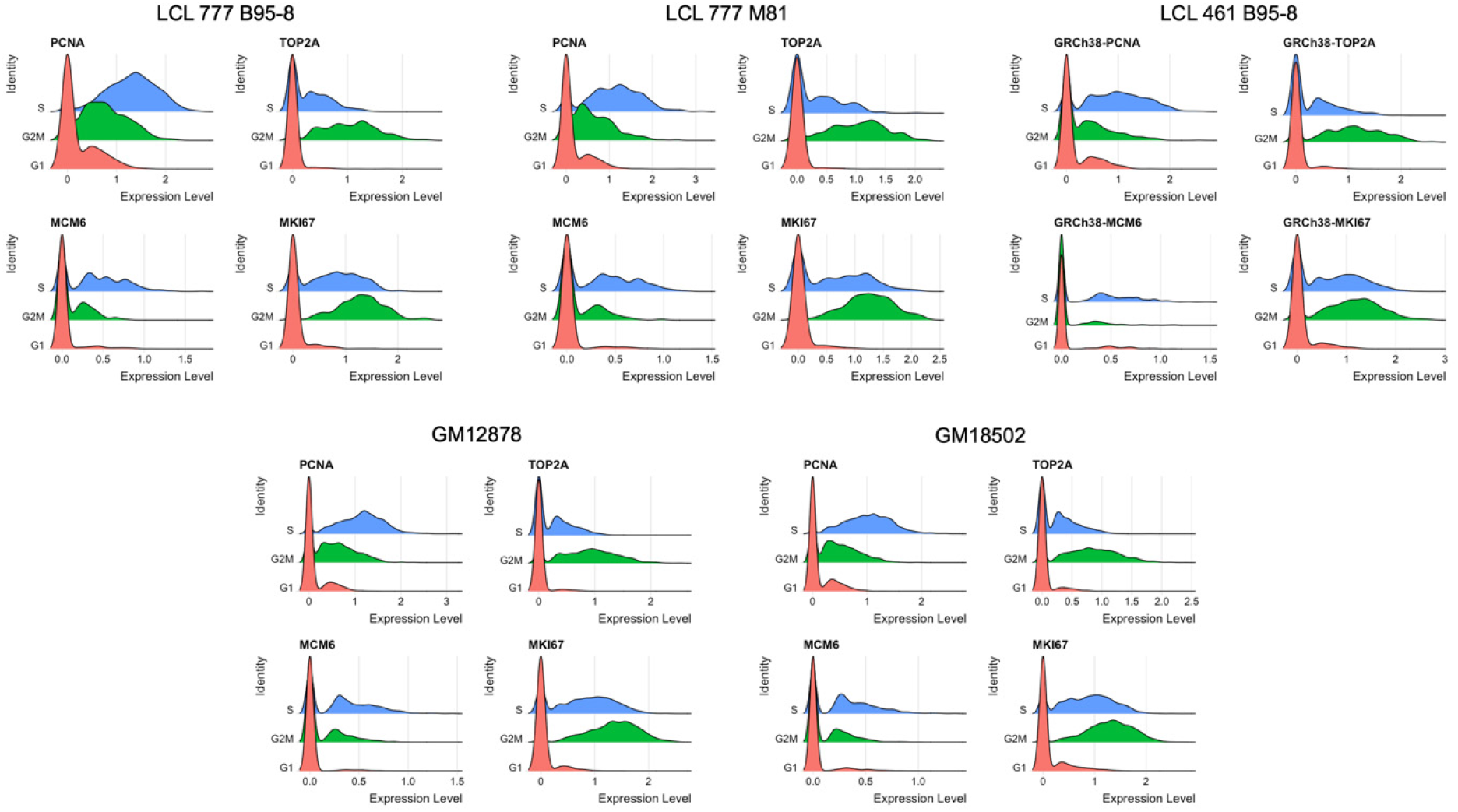
Distributions of representative markers used for cell cycle scoring and regression.

**Figure 1 – figure supplement 4.**
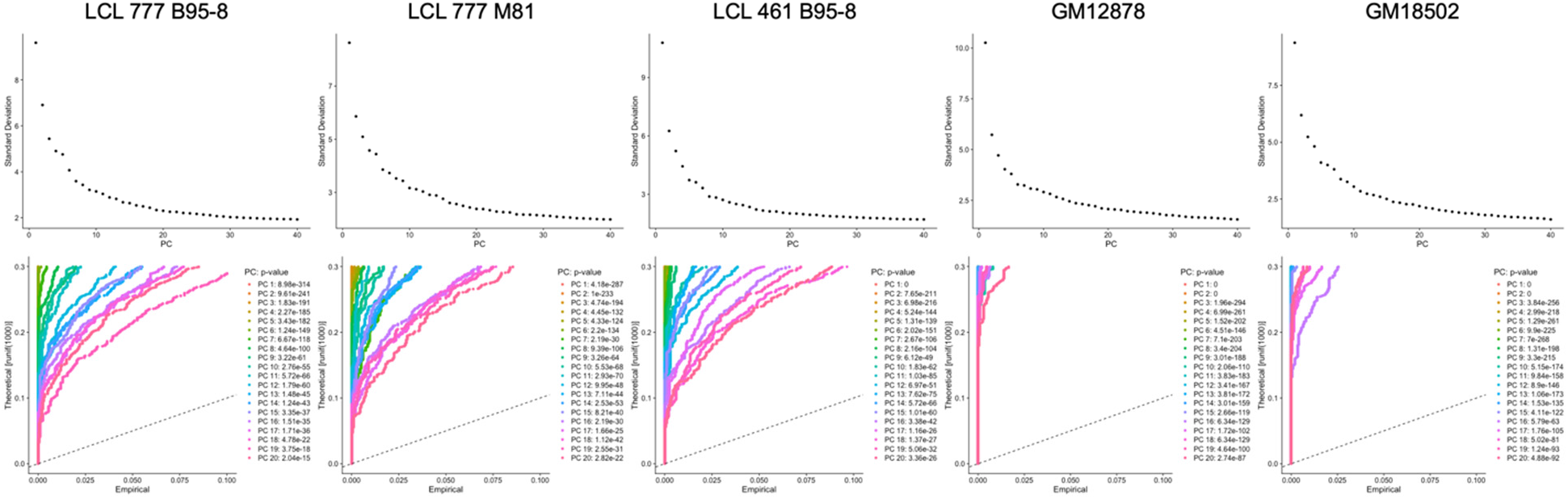
Elbow and Jackstraw plots used for determination of principal components to use for dimensional reduction and clustering.

**Figure 1 – figure supplement 5.**
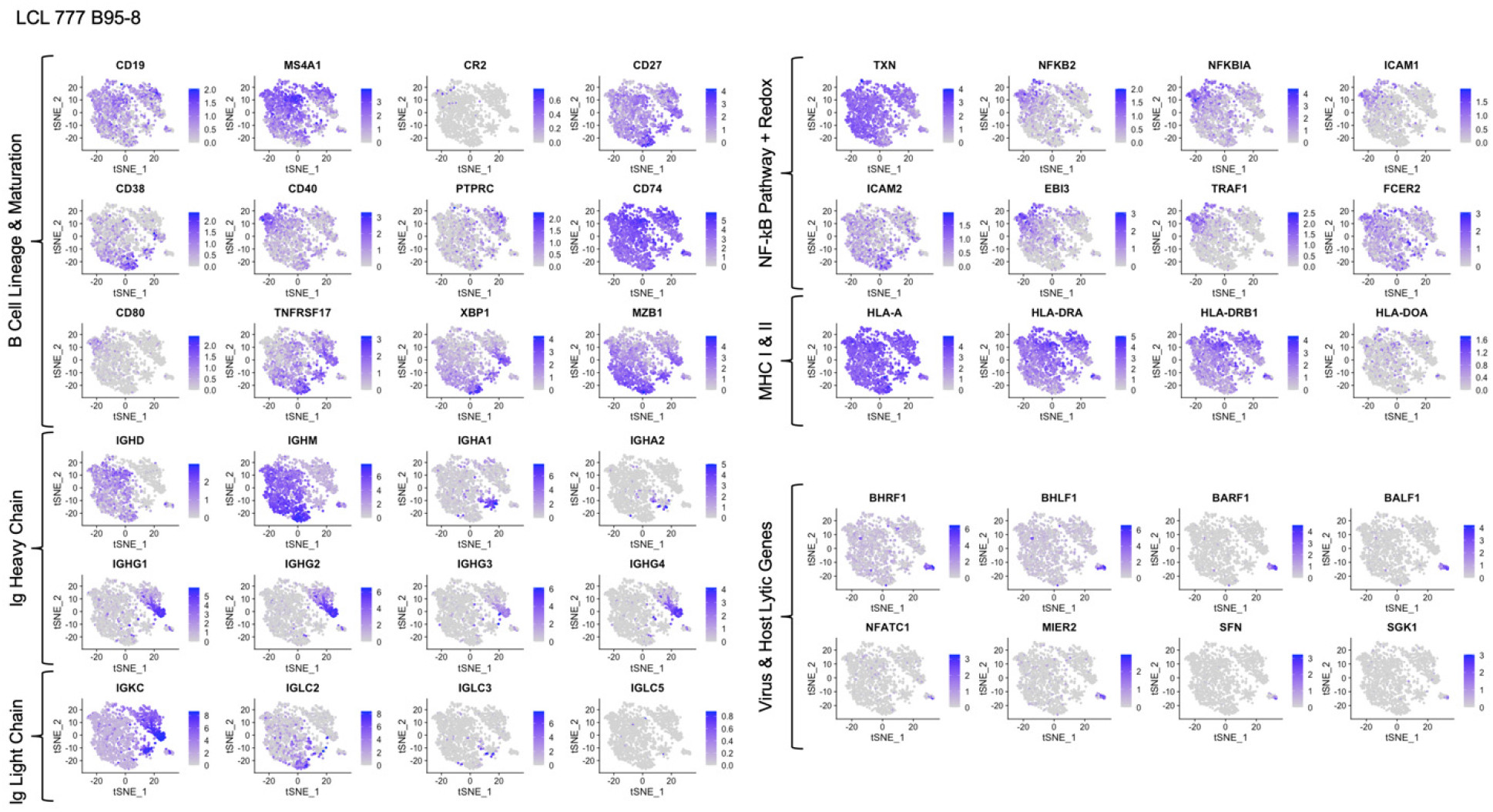
Expression of key genes groups in LCL 777 B95-8.

**Figure 1 – figure supplement 6.**
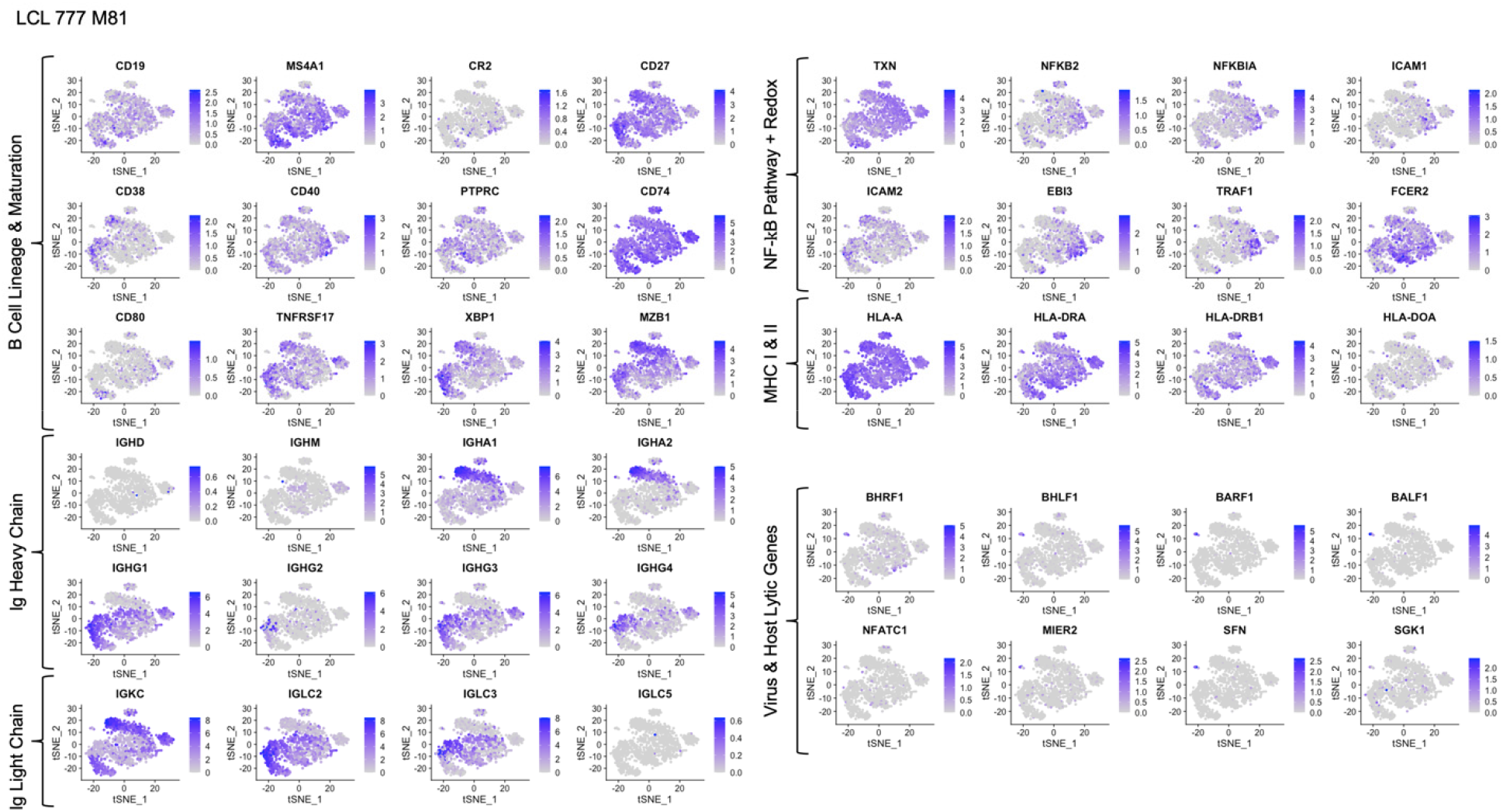
Expression of key genes groups in LCL 777 M81.

**Figure 1 – figure supplement 7.**
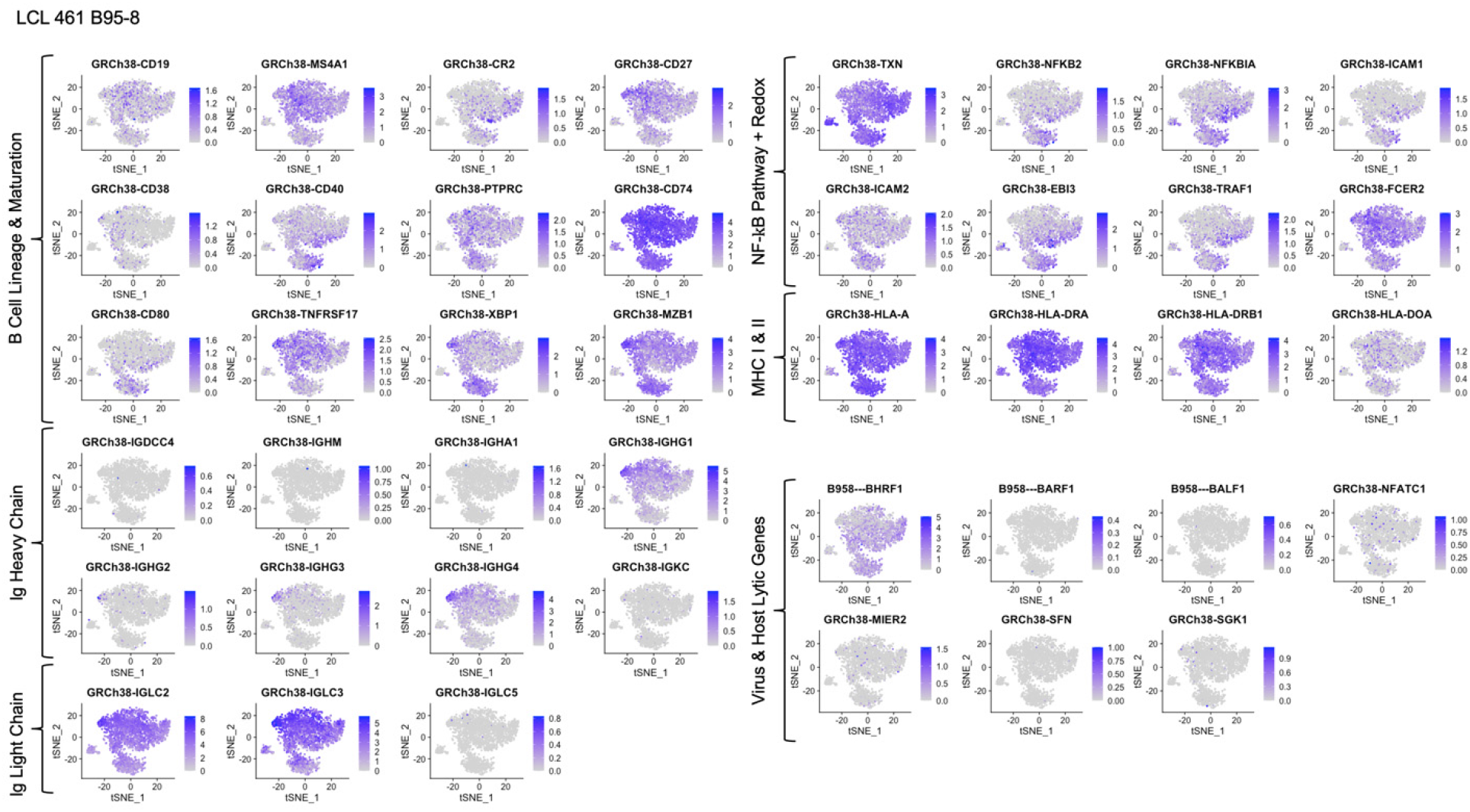
Expression of key genes groups in LCL 461 B95-8.

**Figure 1 – figure supplement 8.**
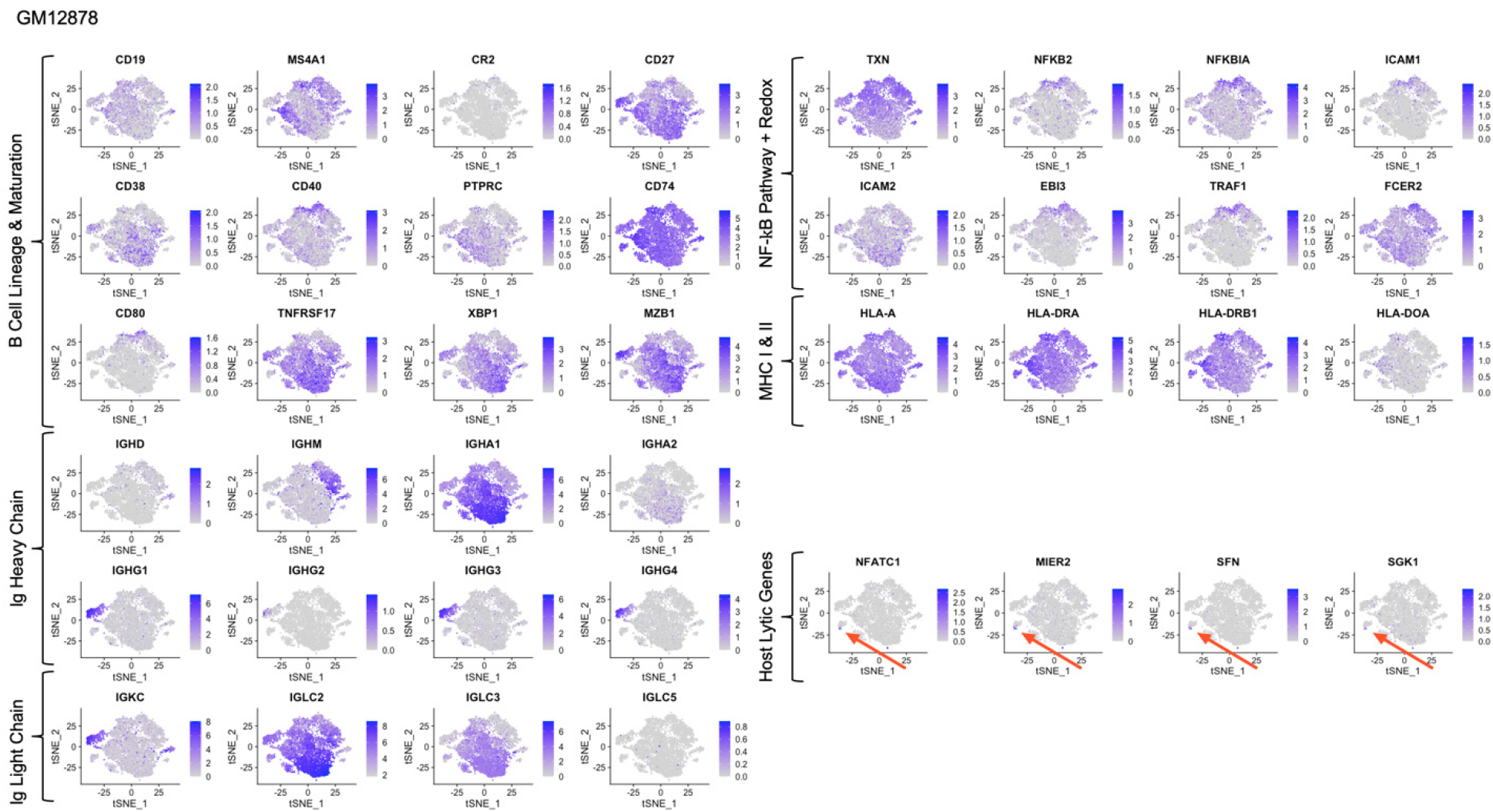
Expression of key genes groups in GM12878.

**Figure 1 – figure supplement 9.**
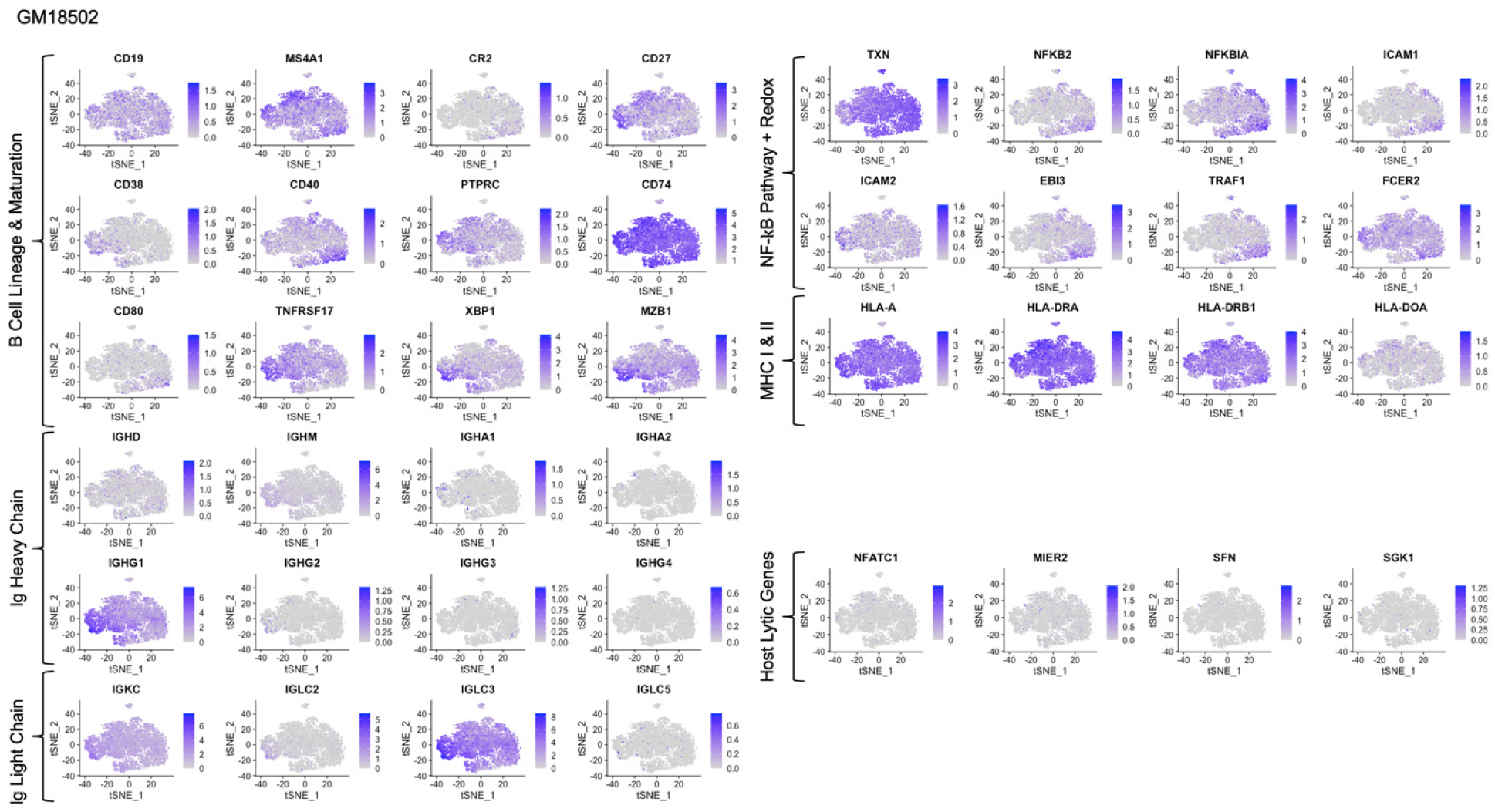
Expression of key genes groups in GM18502.

**Figure 1 – figure supplement 10.**
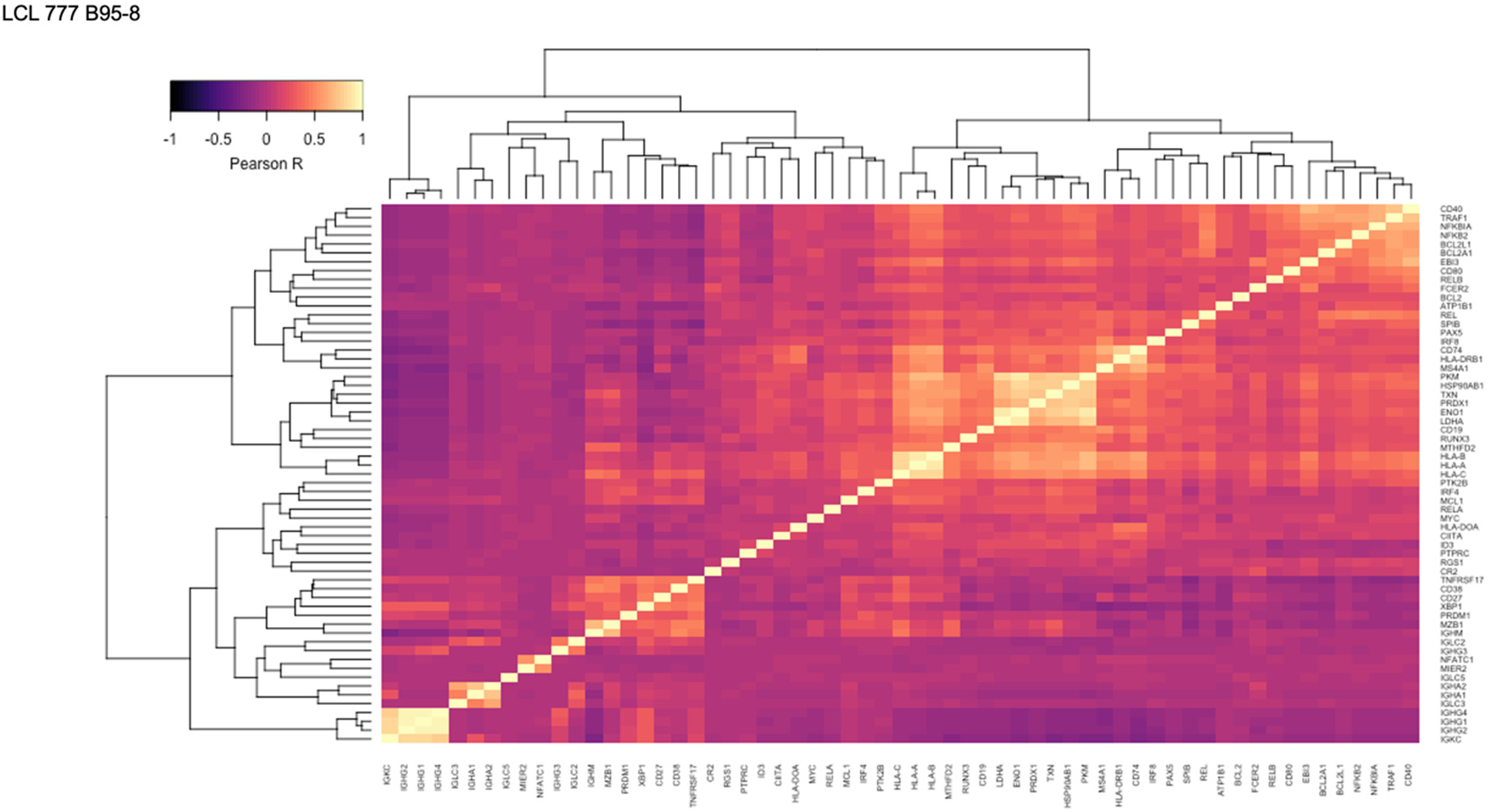
Pairwise Pearson correlation values across key gene groups in LCL 777 B95-8.

**Figure 1 – figure supplement 11.**
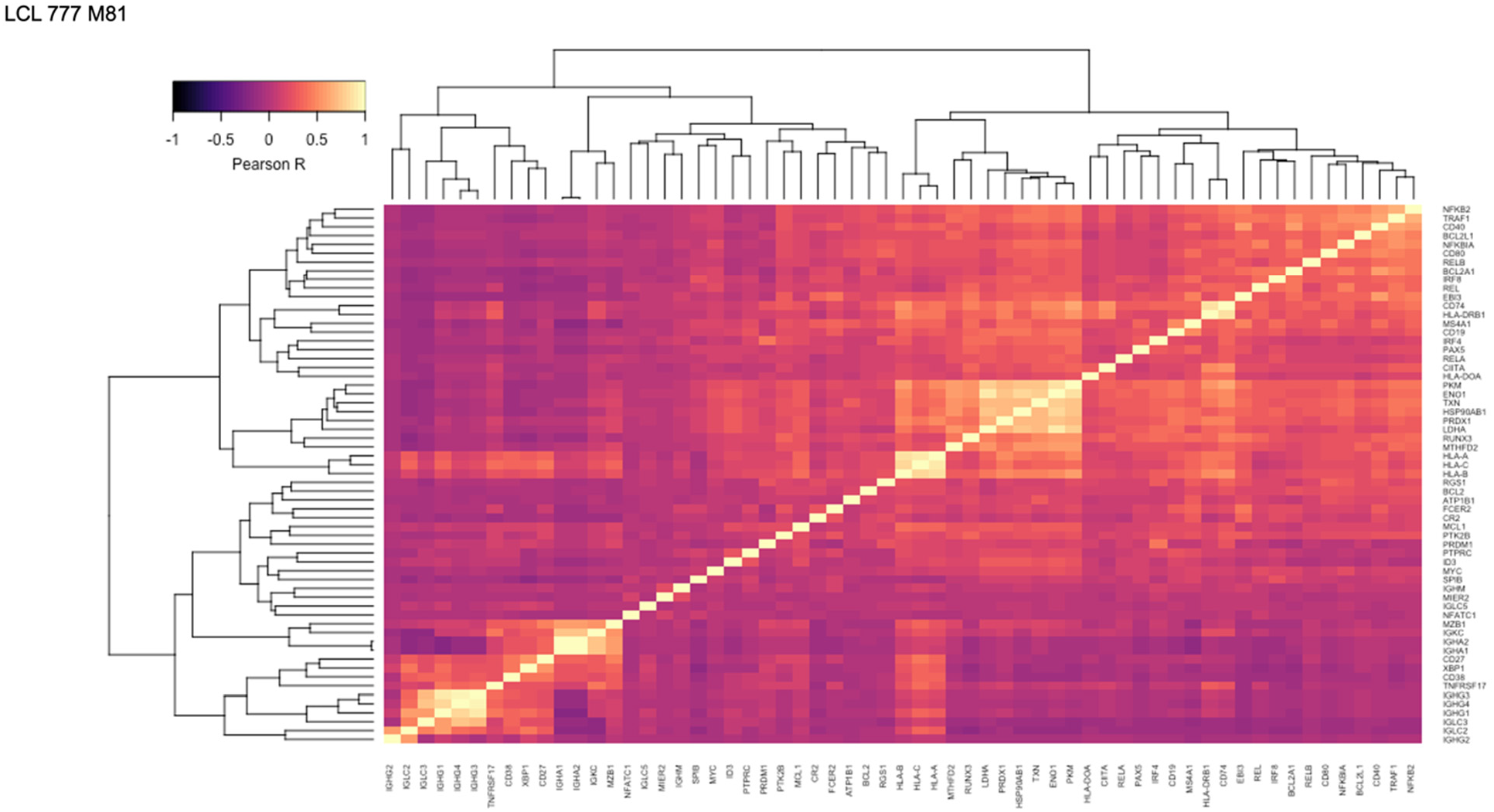
Pairwise Pearson correlation values across key gene groups in LCL 777 M81.

**Figure 1 – figure supplement 12.**
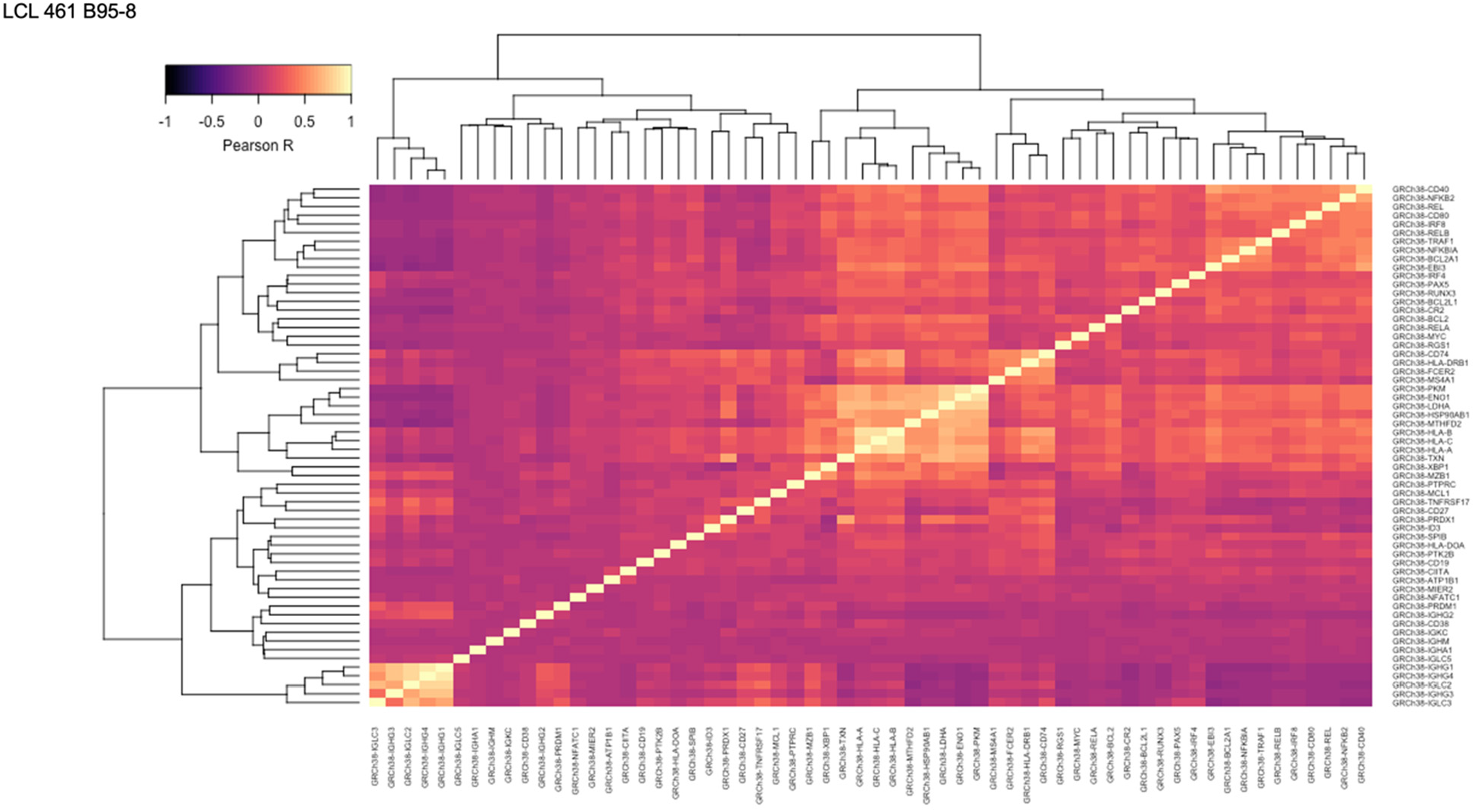
Pairwise Pearson correlation values across key gene groups in LCL 461 B95-8.

**Figure 1 – figure supplement 13.**
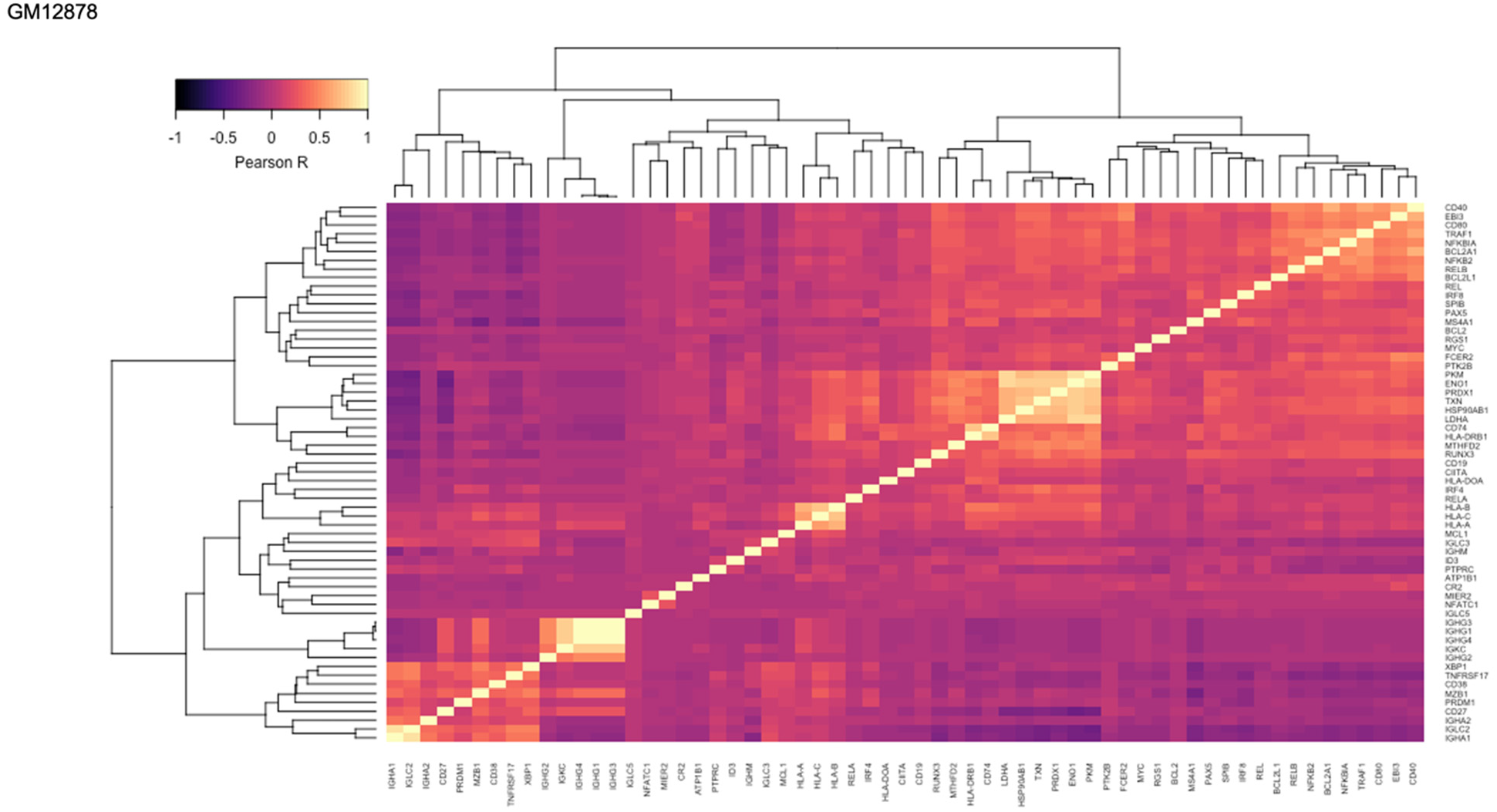
Pairwise Pearson correlation values across key gene groups in GM12878.

**Figure 1 – figure supplement 14.**
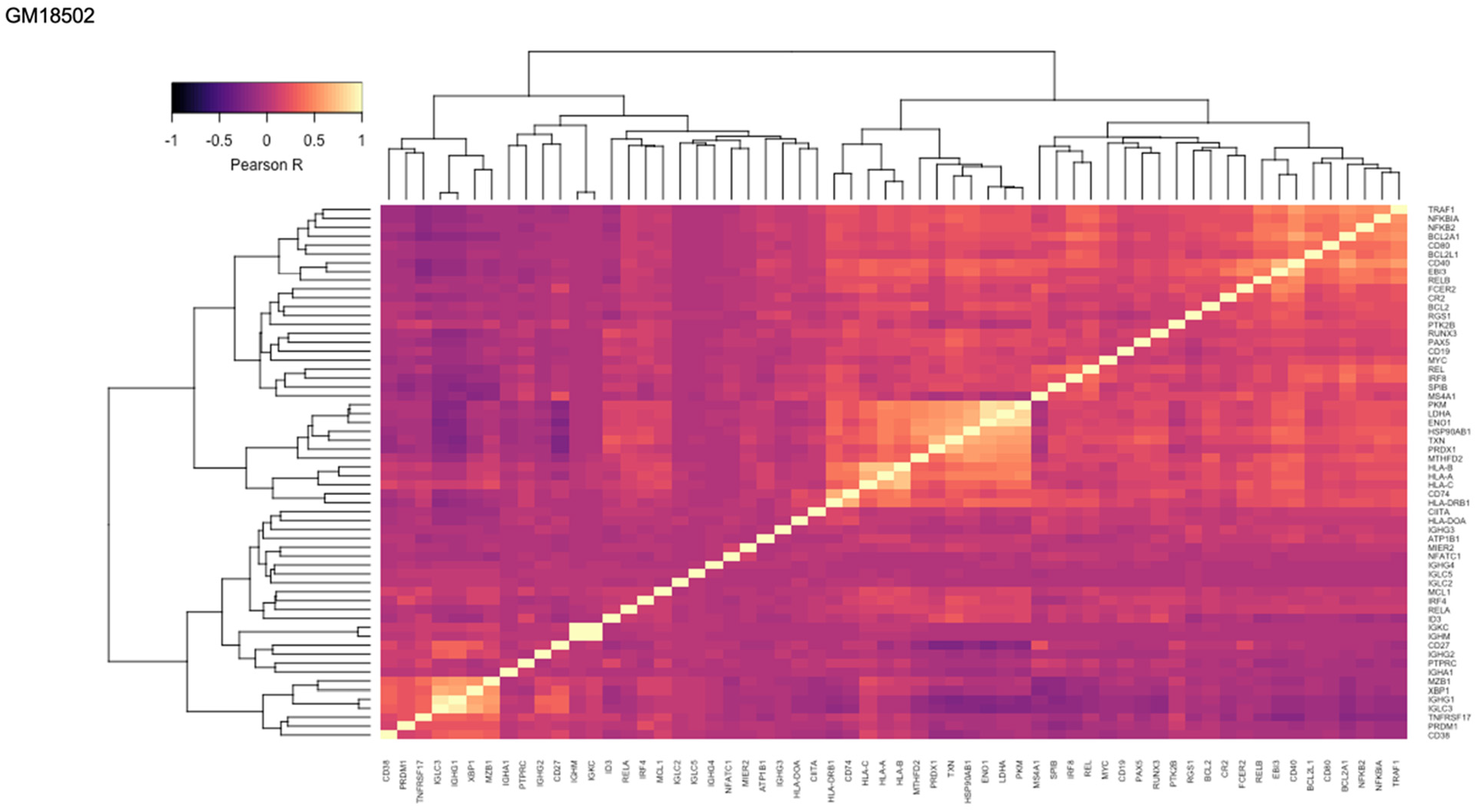
Pairwise Pearson correlation values across key gene groups in GM18502.

**Figure 1 – figure supplement 15.**
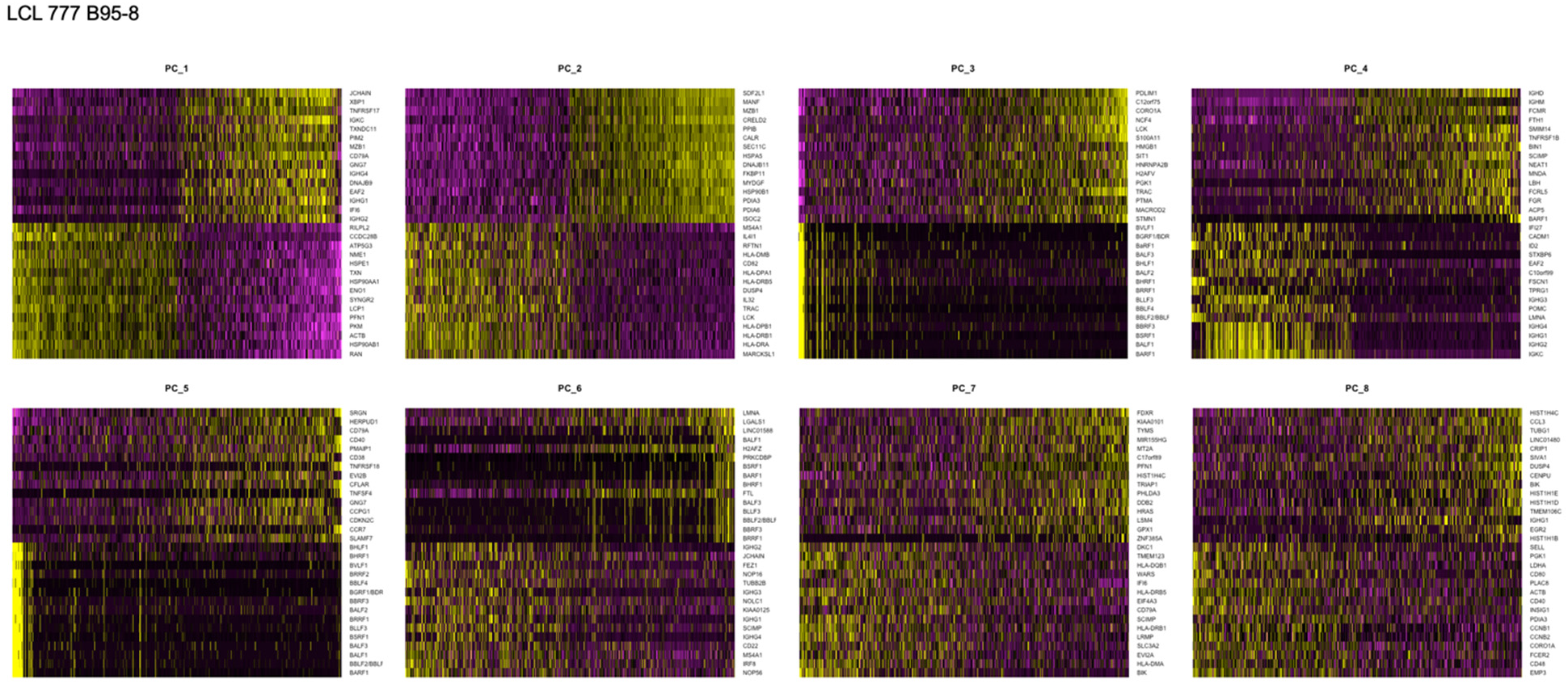
Principal component (PCs 1-8) heatmaps for LCL 777 B95-8.

**Figure 1 – figure supplement 16.**
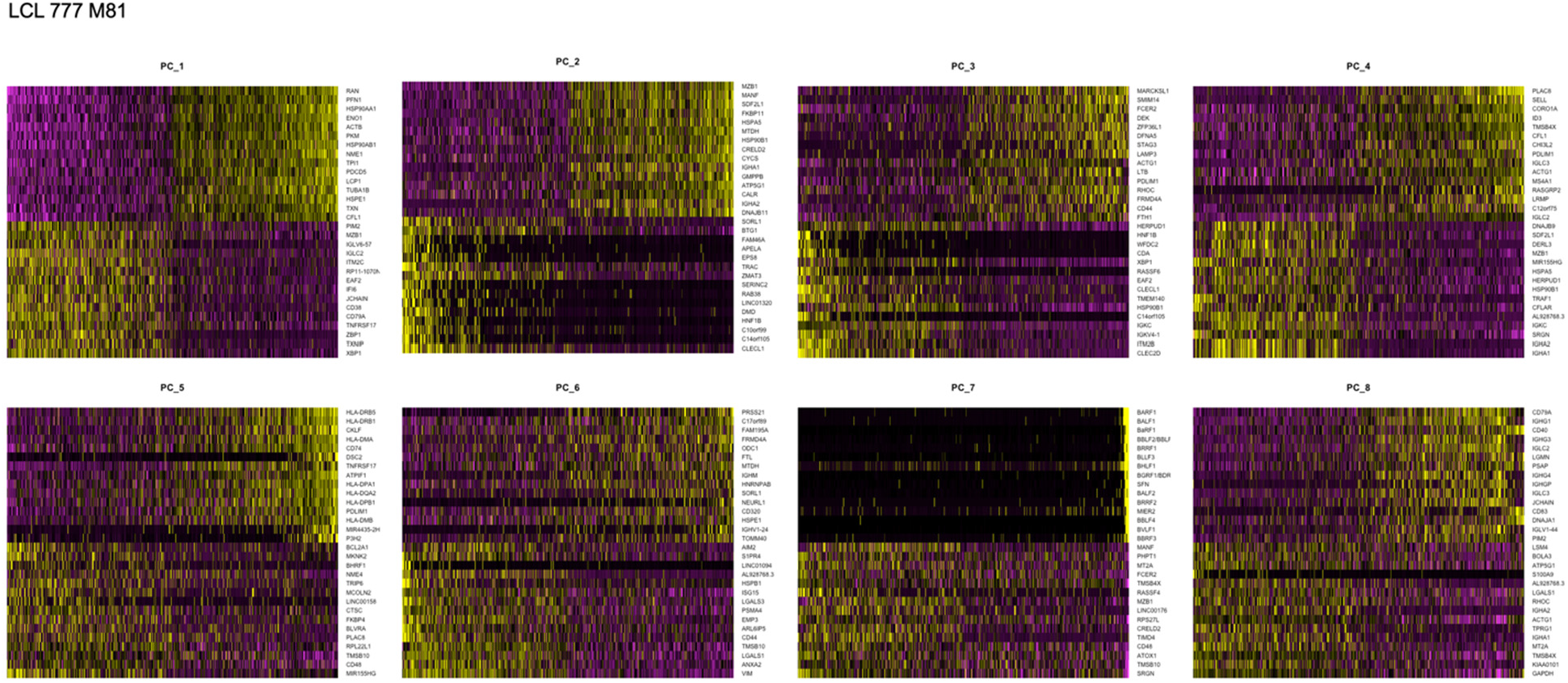
Principal component (PCs 1-8) heatmaps for LCL 777 M81.

**Figure 1 – figure supplement 17.**
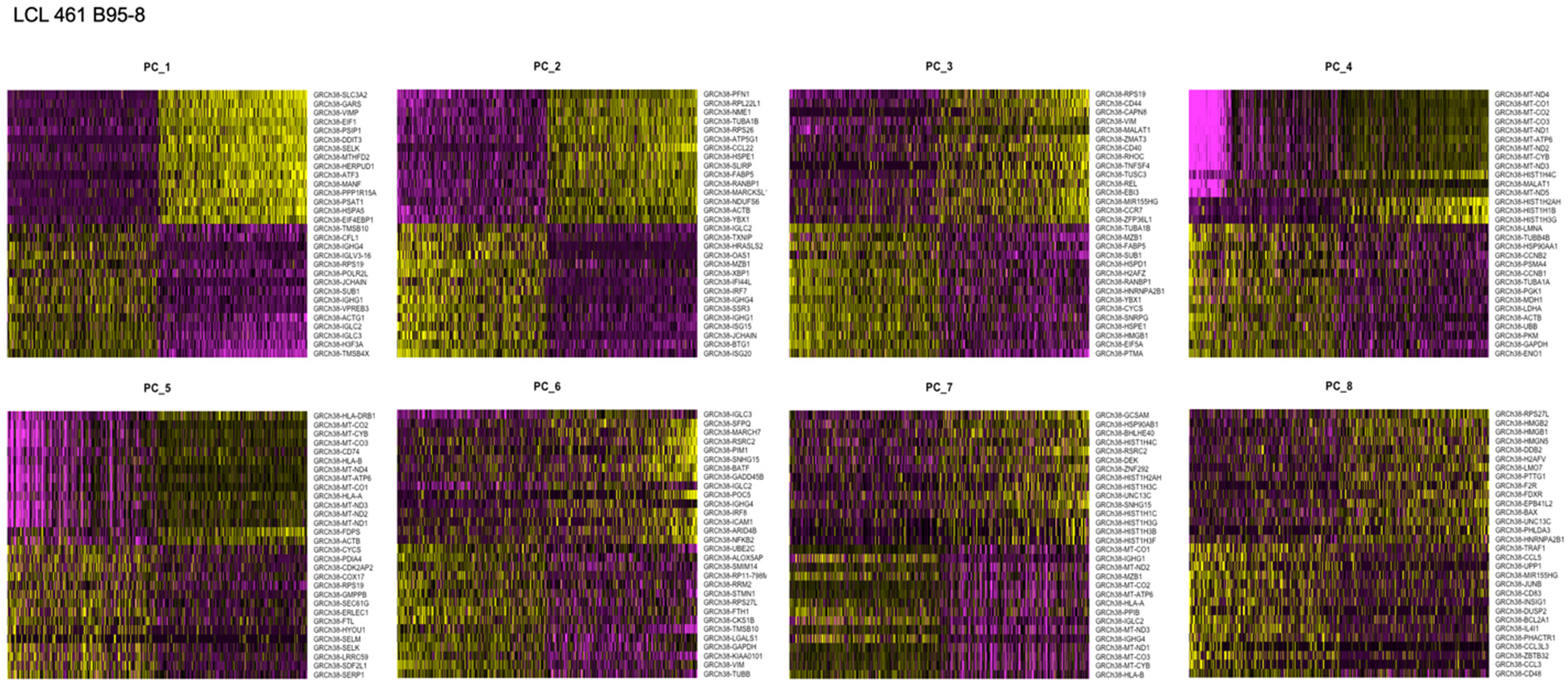
Principal component (PCs 1-8) heatmaps for LCL 461 B95-8.

**Figure 1 – figure supplement 18.**
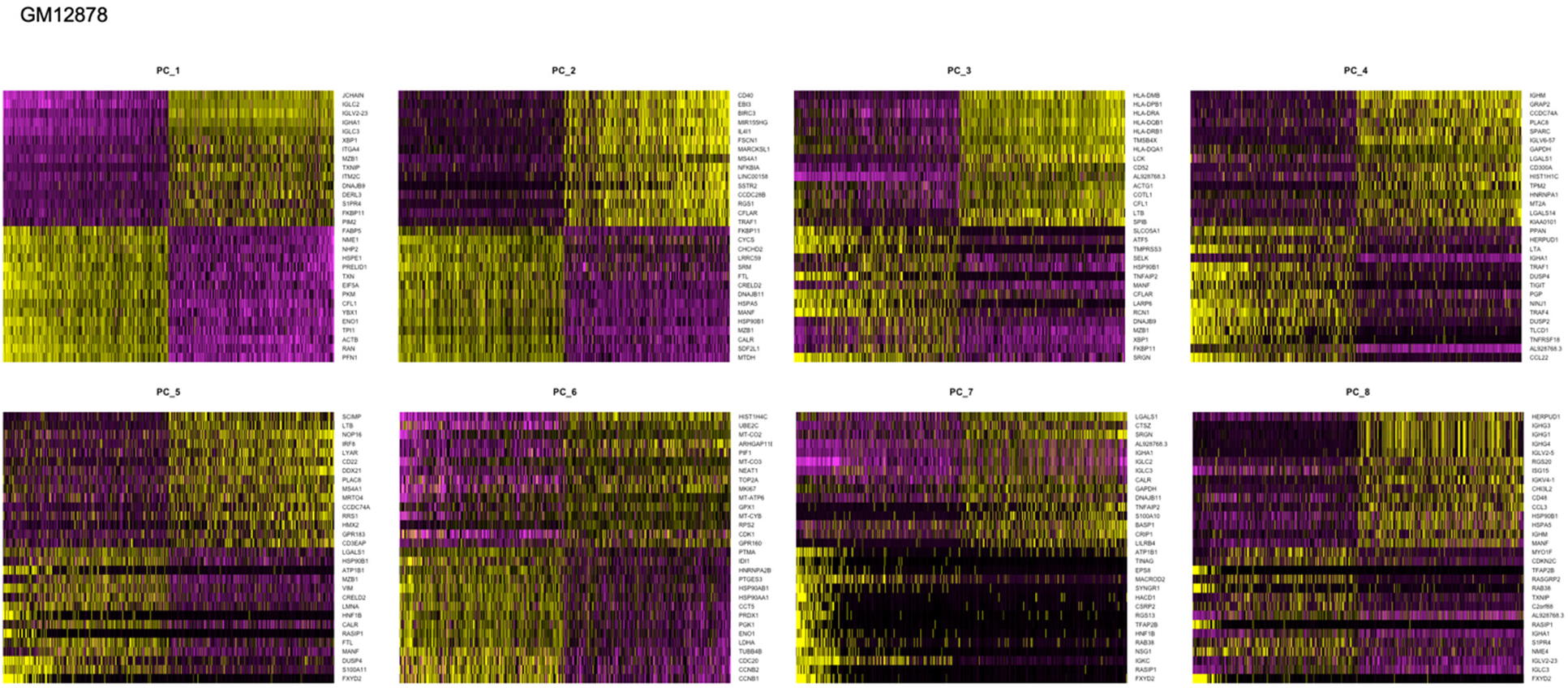
Principal component (PCs 1-8) heatmaps for GM12878.

**Figure 1 – figure supplement 19.**
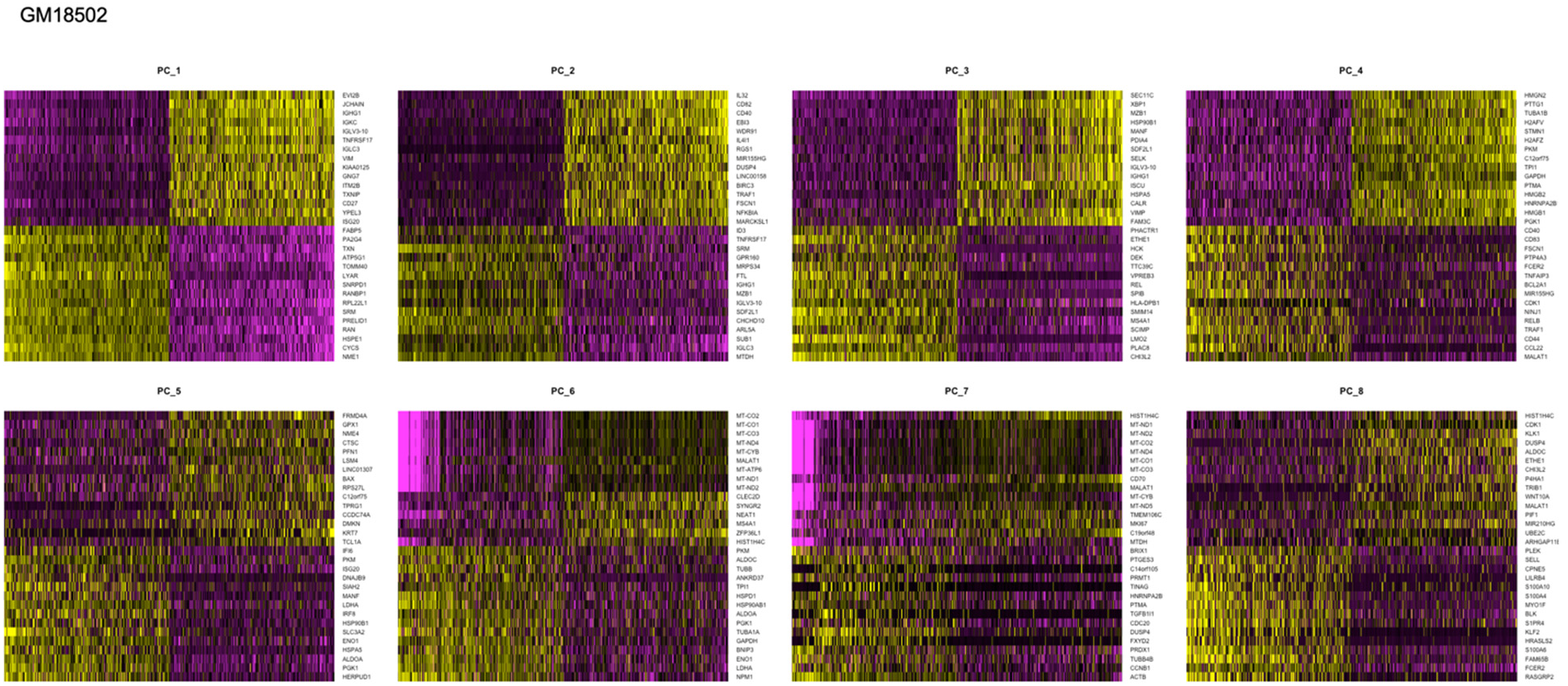
Principal component (PCs 1-8) heatmaps for GM18502.

**Figure 2 – figure supplement 1.**
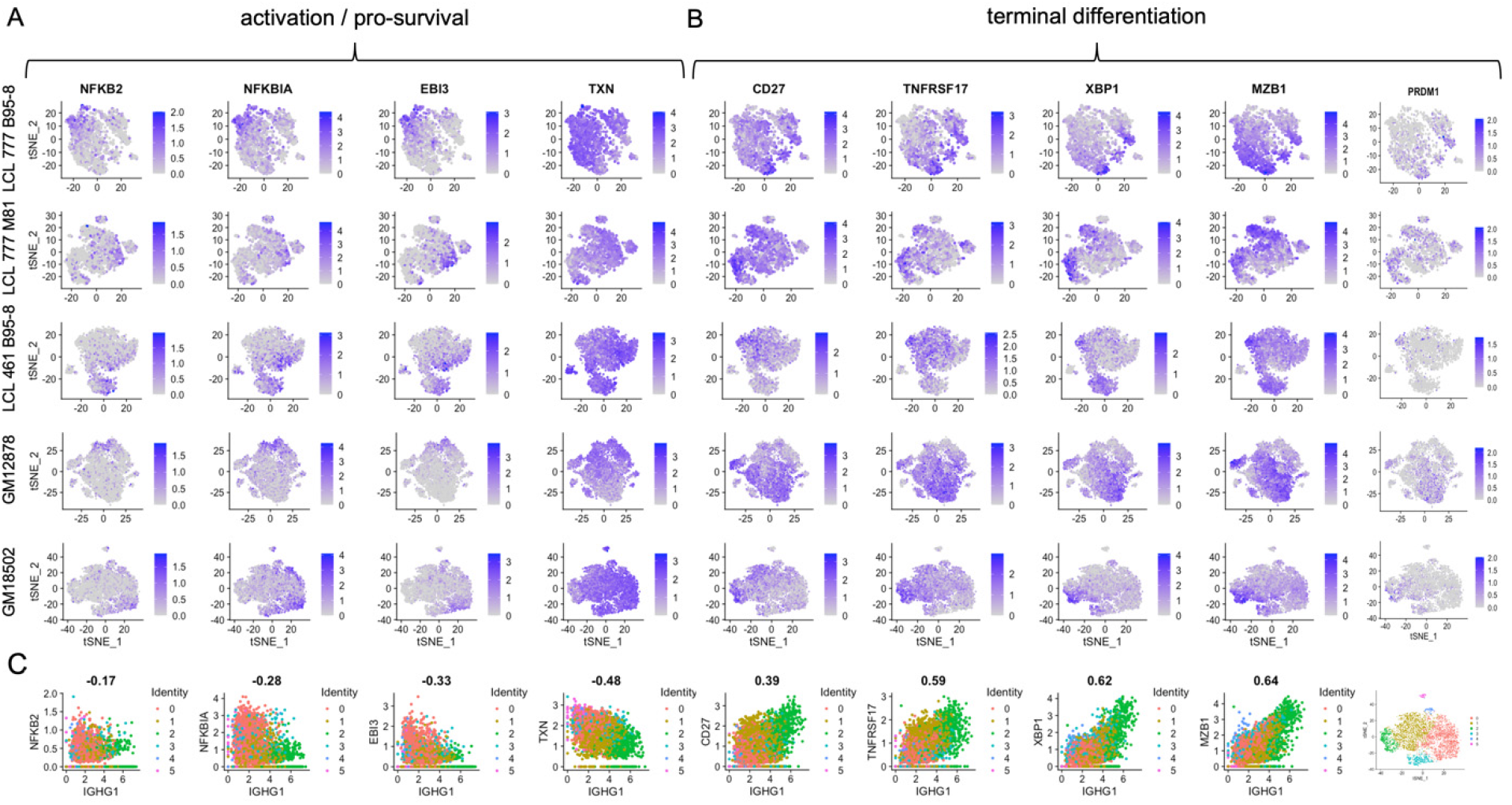
Expression of individual genes within activation and differentiation gene sets. **(A)** Expression of genes involved in activation / pro-survival of B cells. **(B)** Expression of genes involved in B cell differentiation. **(C)** Correlation of each activation and differentiation gene with IgG1 expression for GM18502 (a sample with a single isotype class).

**Figure 2 – figure supplement 2.**
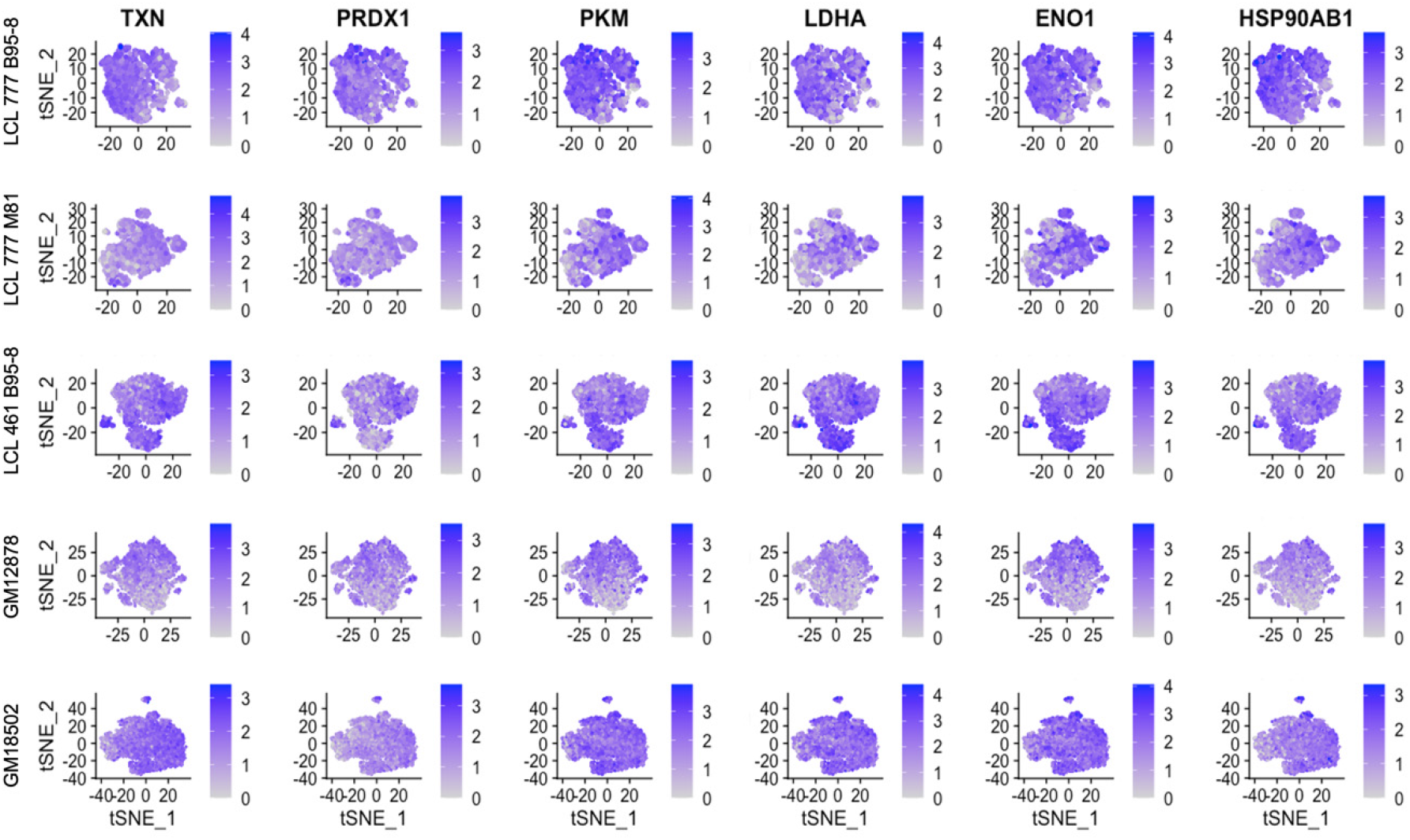
Expression of metabolic and oxidative stress genes.

**Figure 2 – figure supplement 3.**
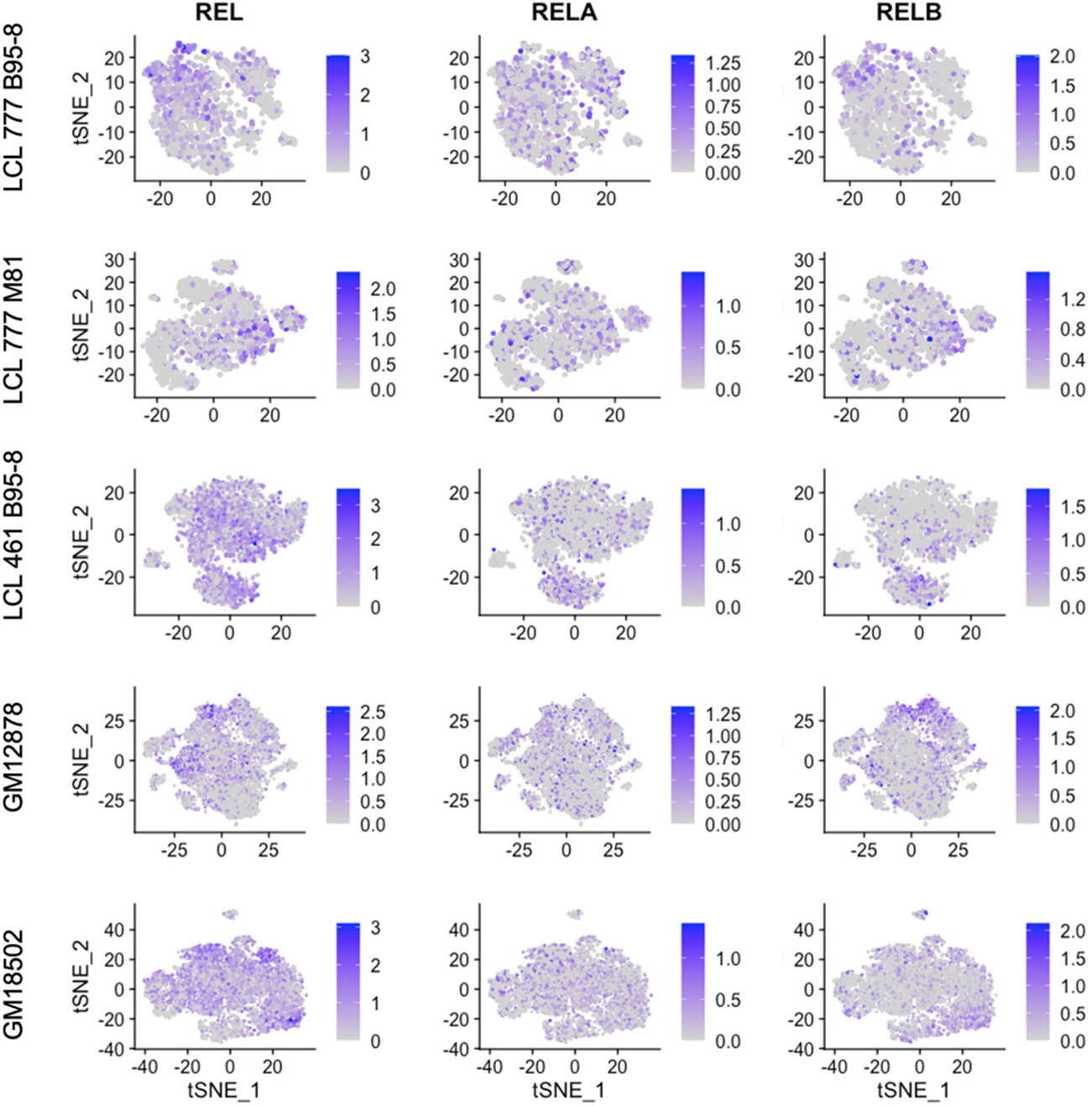
Expression of NF-kB subunits c-REL, RELA, and RELB.

**Figure 2 – figure supplement 4.**
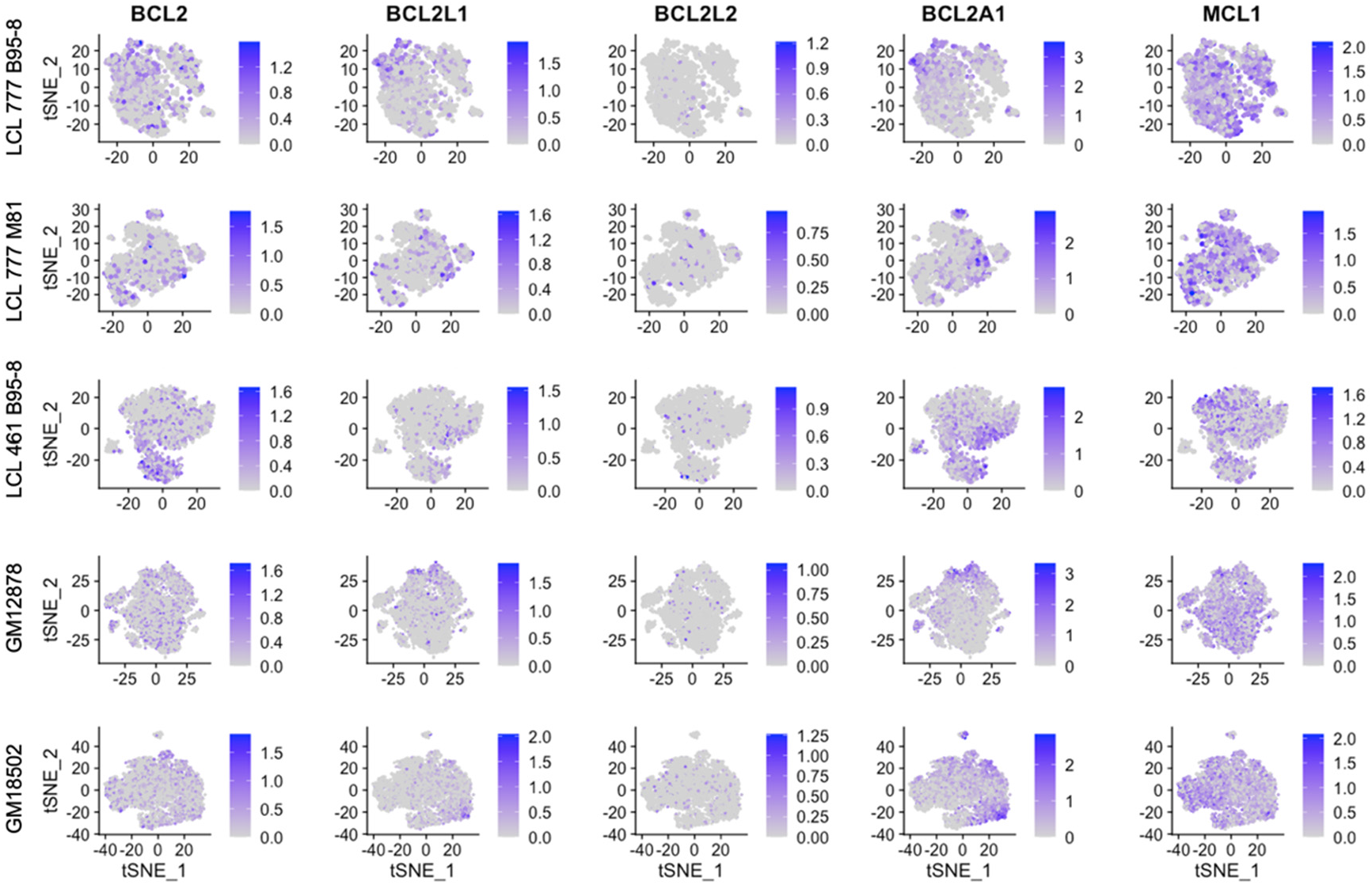
Expression of BCL family genes across LCL samples.

**Figure 2 – figure supplement 5.**
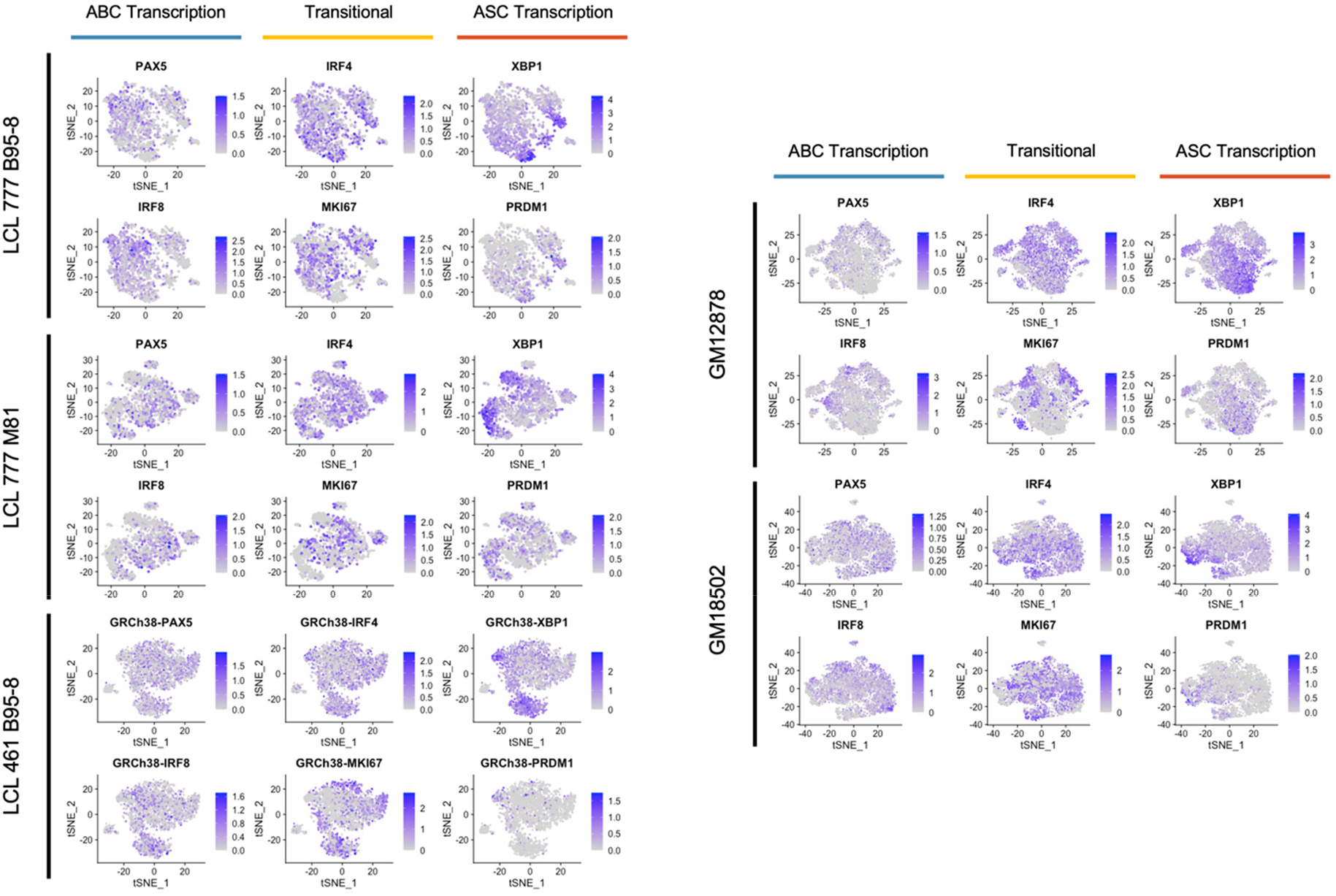
Expression trends in key transcriptional regulators controlling activated B cell (ABC) and antibody-secreting cell (ASC) phenotypes.

**Figure 2 – figure supplement 6.**
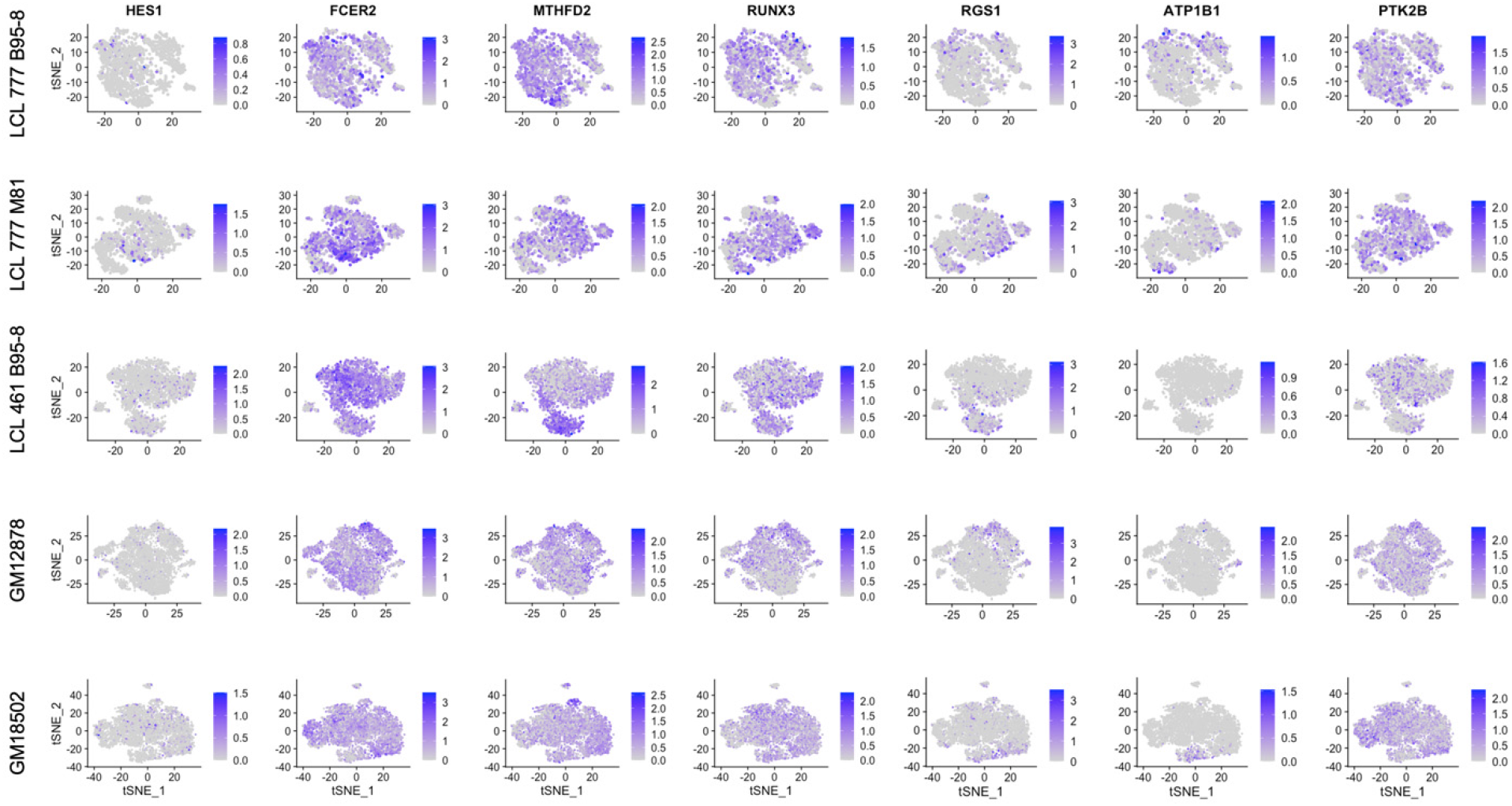
Expression of host targets upregulated by EBNA2.

**Figure 2 – figure supplement 7.**
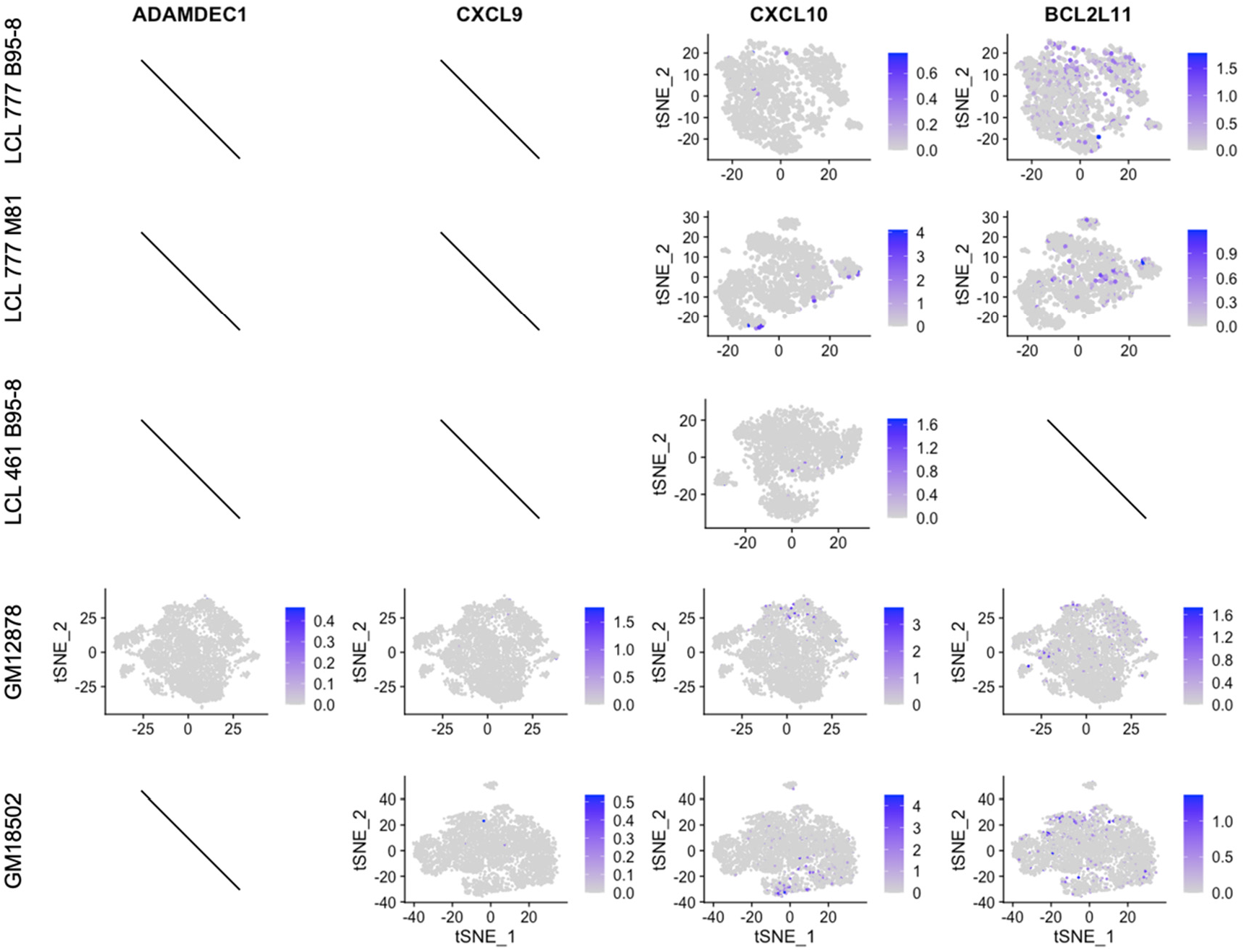
Expression of host targets repressed by EBNA3.

**Figure 4 – figure supplement 1.**
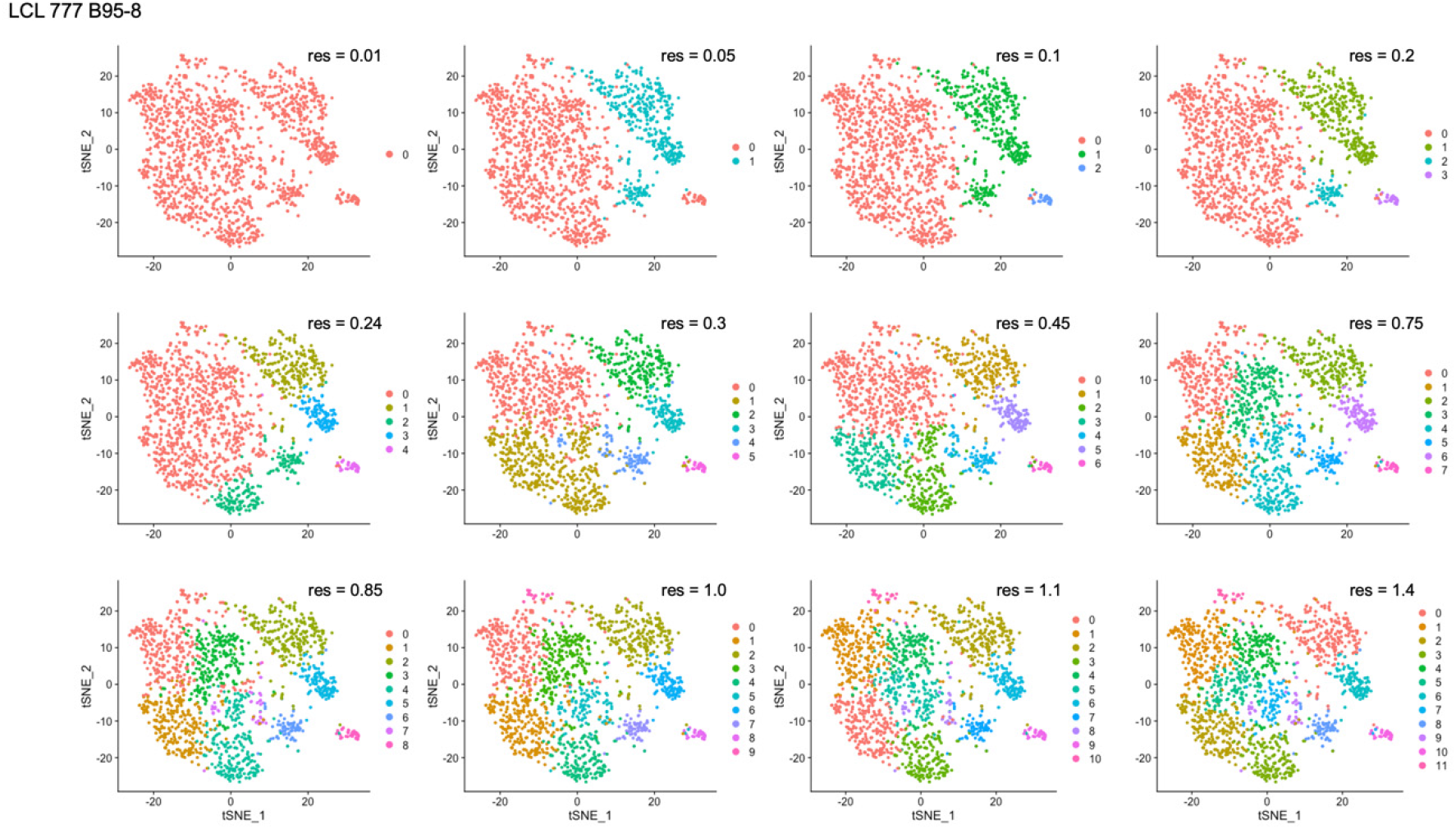
Clustering resolution screens for LCL 777 B95-8.

**Figure 4 – figure supplement 2.**
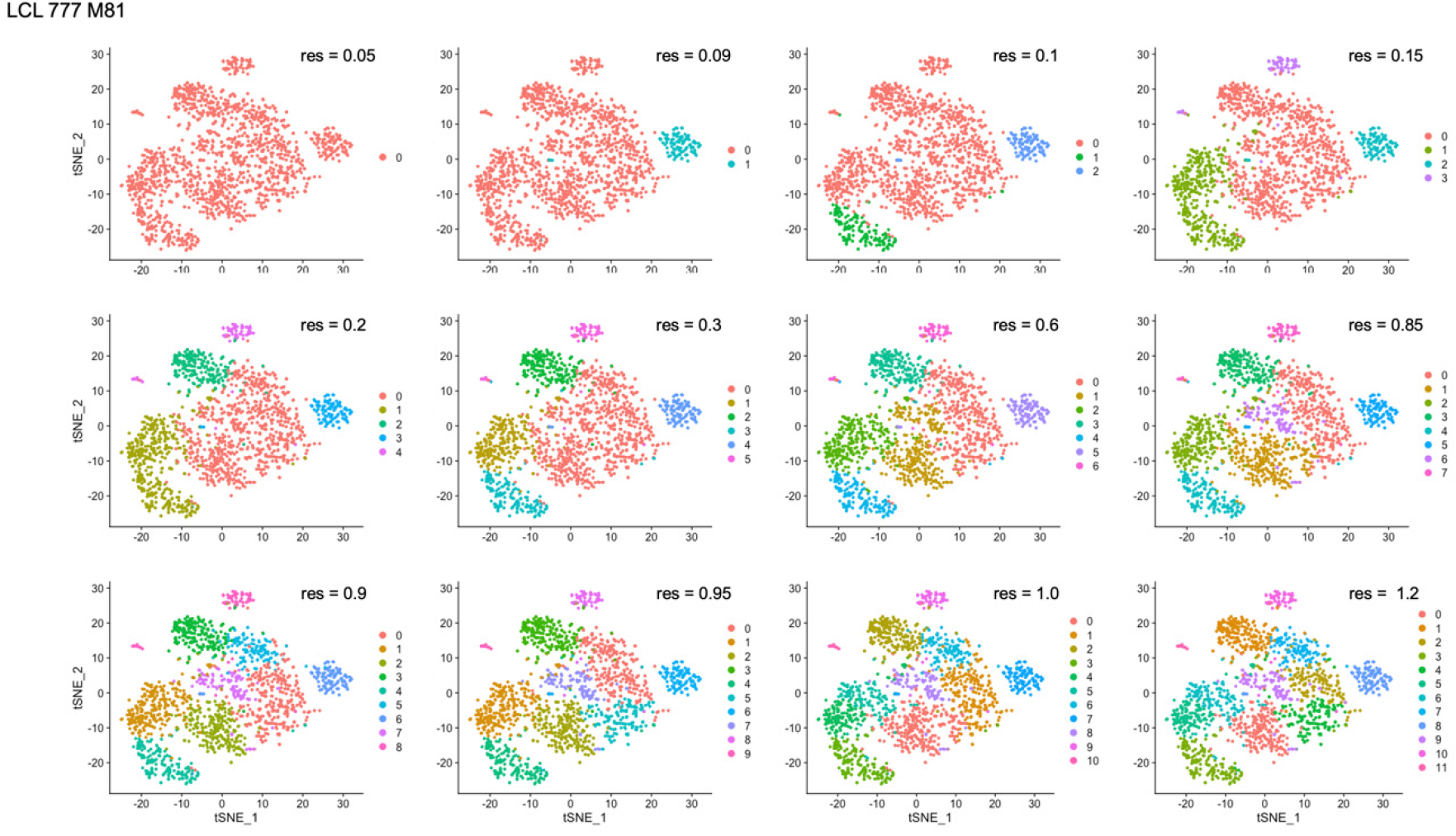
Clustering resolution screens for LCL 777 M81.

**Figure 4 – figure supplement 3.**
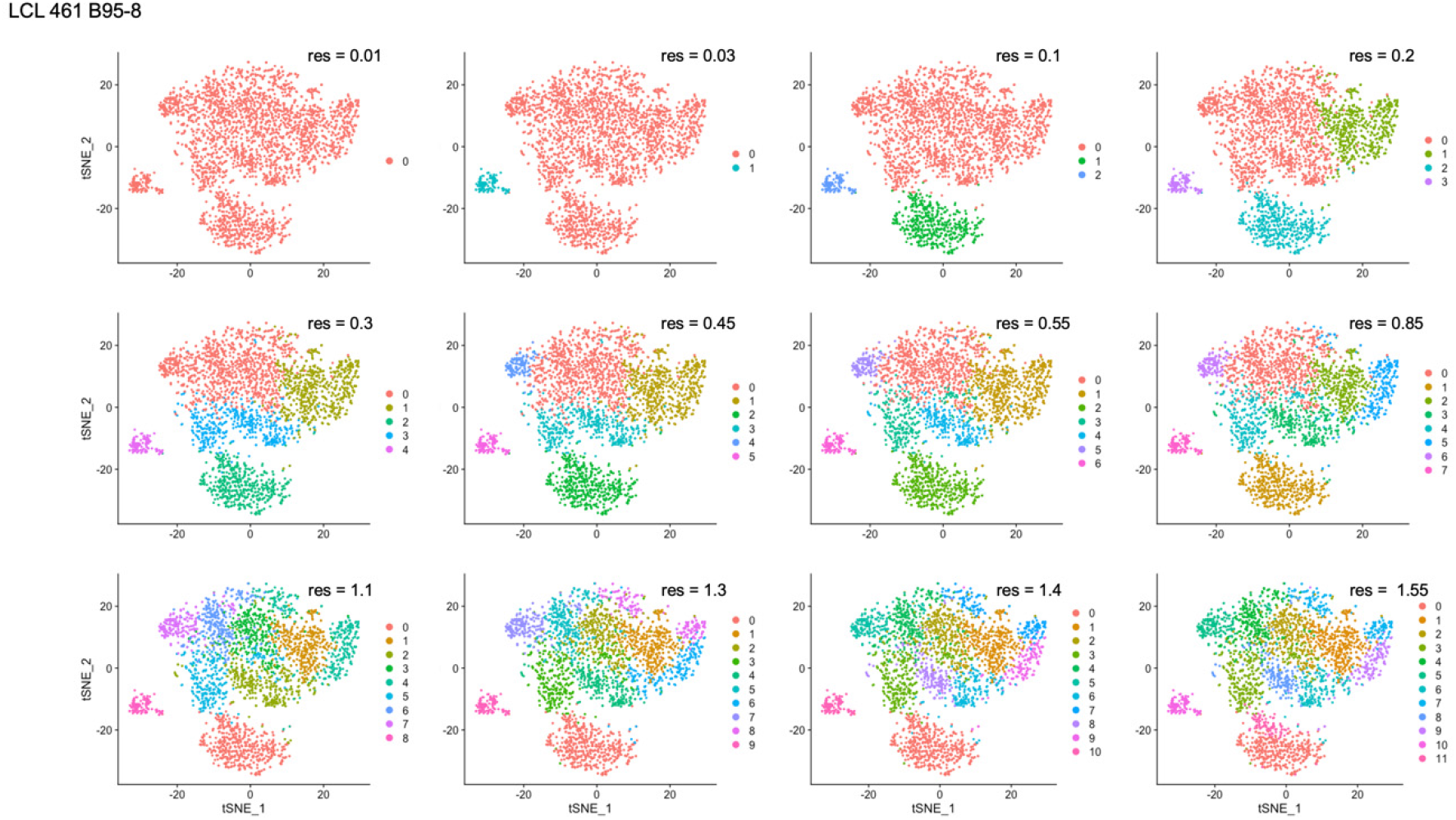
Clustering resolution screens for LCL 461 B95-8.

**Figure 4 – figure supplement 4.**
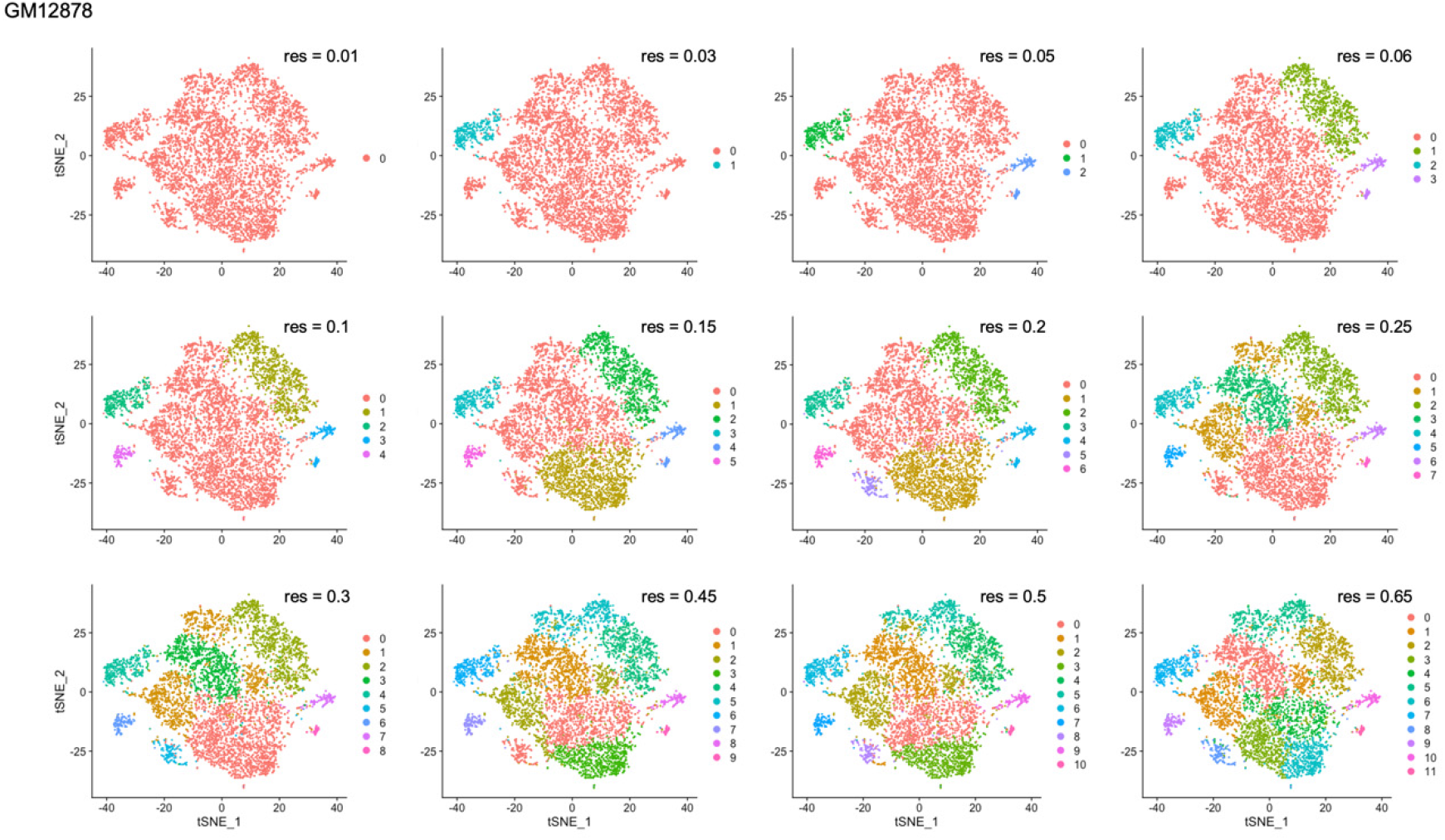
Clustering resolution screens for GM12878.

**Figure 4 – figure supplement 5.**
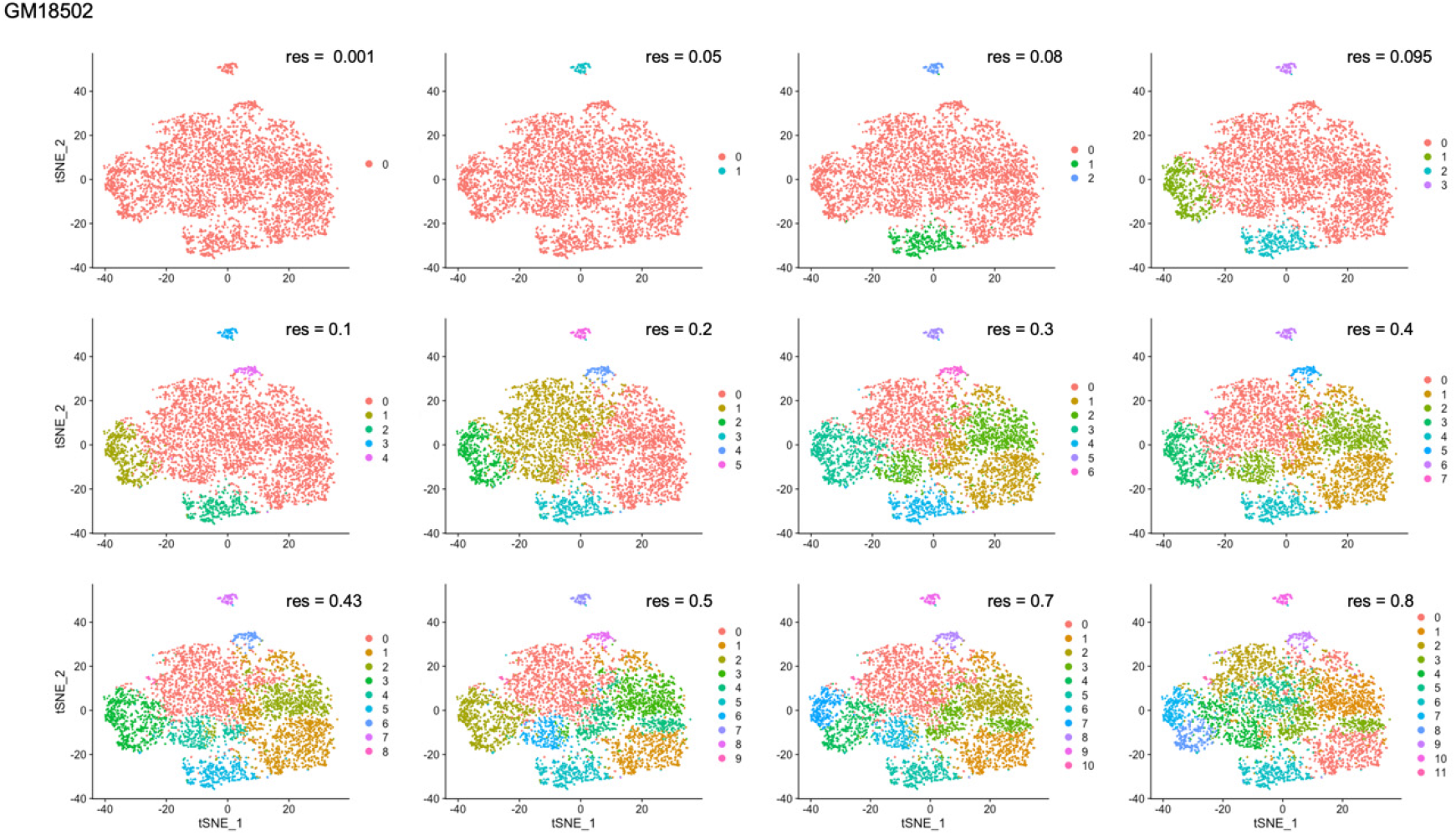
Clustering resolution screens for GM18502.

**Figure 4 – figure supplement 6.**
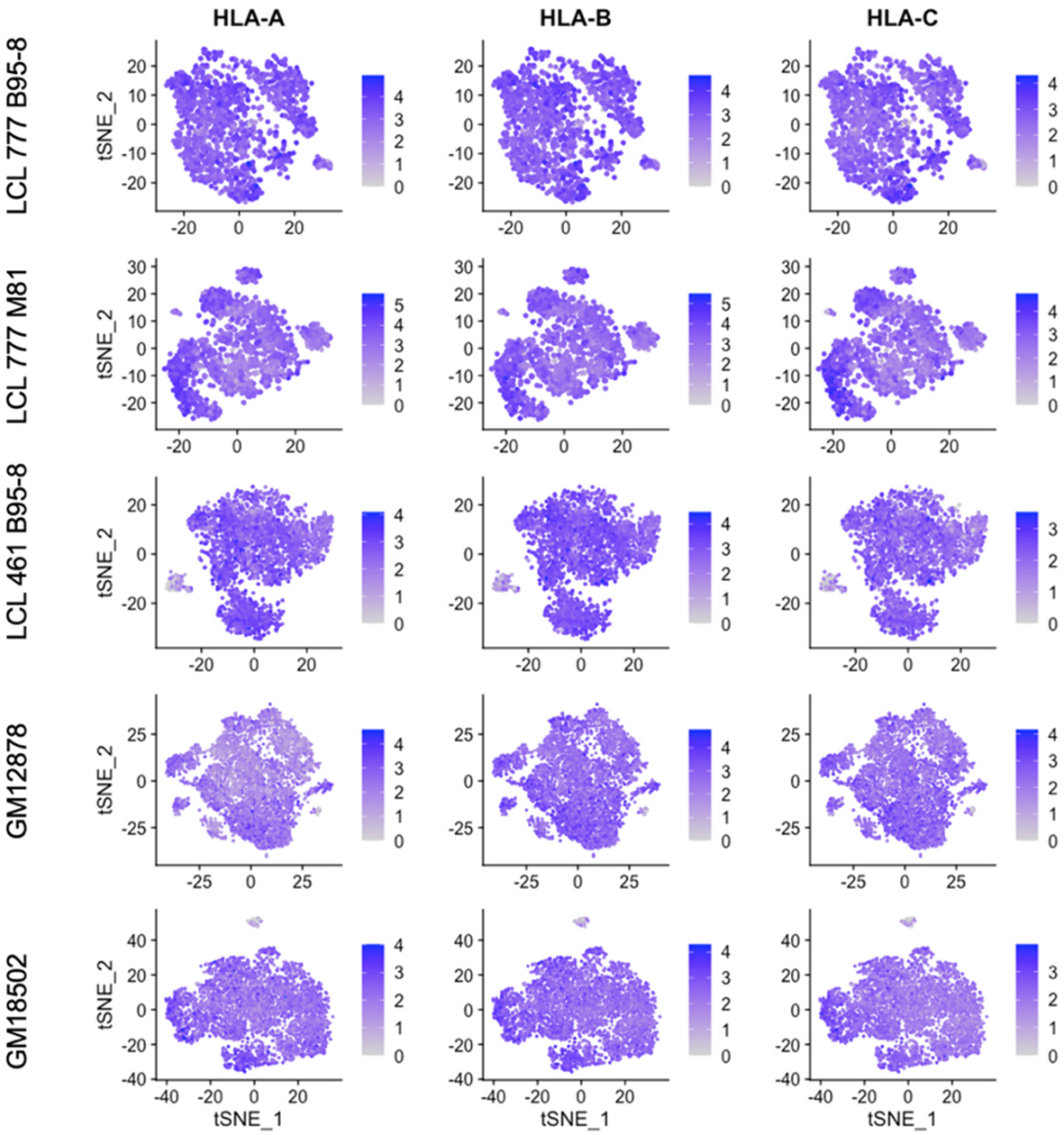
Expression of MHC class I genes HLA-A, HLA-B, and HLA-C.

**Figure 4 – figure supplement 7.**
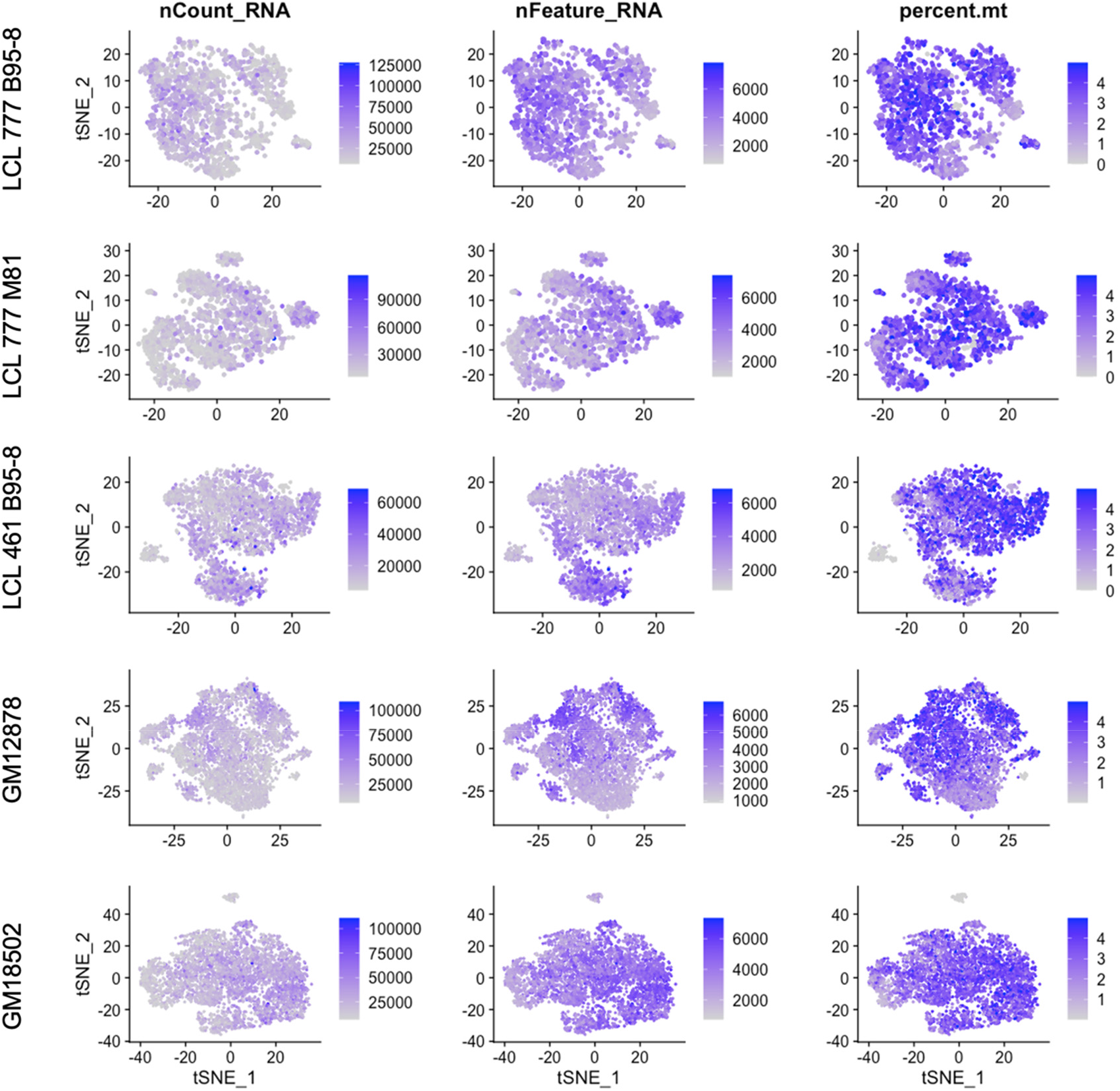
Total RNA counts, unique feature, and mitochondrial percentage distributions across LCL samples.

